# RNA binding is essential for NONO condensates to modulate pre-mRNA processing of super enhancer-associated genes in neuroblastoma

**DOI:** 10.1101/2022.02.28.482217

**Authors:** Song Zhang, Jack Cooper, Yee Seng Chong, Alina Naveed, Chelsea Mayoh, Nisitha Jayatilleke, Tao Liu, Sebastian Amos, Simon Kobelke, Andrew C Marshall, Oliver Meers, Yu Suk Choi, Charles S Bond, Archa H Fox

## Abstract

High-risk neuroblastoma patients have poor survival rates and require better therapeutic options. High expression of a multifunctional DNA and RNA binding protein, NONO, in neuroblastoma is associated with poor patient outcome, however there is little understanding of the mechanism of NONO-dependent oncogenic gene regulatory activity in neuroblastoma. Here, we used cell imaging, biophysical and molecular analysis to reveal complex NONO-dependent regulation of gene expression, finding that NONO forms RNA- and DNA-tethered phase-separated condensates throughout the nucleus. CLIP analyses show that NONO mainly binds to the 5’ end of pre-mRNAs and modulates pre-mRNA processing, dependent on its RNA binding activity. NONO preferentially regulates super enhancer-associated genes, including HAND2 and GATA2. In the absence of functional NONO-RNA condensates, inefficient pre-mRNA processing at these loci leads to decreased expression of HAND2 and GATA2. Thus, future development of agents that target RNA binding activity of NONO may have therapeutic potential in this cancer context.

## Introduction

Neuroblastoma is derived from neural crest cells in the sympathetic nervous system and is the most common extracranial solid cancer in children (Janoueix-Lerosey, Schleiermacher et al., 2010, Munzer, Menegaux et al., 2008). Whilst suitable treatments for low-risk patients exist, high-risk neuroblastoma patients have poor survival rates and a paucity of therapeutic options. High expression of the gene regulatory protein NONO (Non-POU Domain Containing Octamer Binding) in neuroblastoma is associated with poor patient survival, suggesting this could be a potential therapeutic target (Liu, Erriquez et al., 2014). However, beyond knowledge of NONO binding to one long non-coding RNA (lncRNA) in neuroblastoma, LncMycnUS, there is no further insight into NONO mechanistic activity in this cancer. NONO is a multifunctional protein with various roles in genome maintenance and gene regulation at the transcriptional and post-transcriptional levels, including transcription initiation, elongation and termination, pre-mRNA processing and splicing, and nuclear retention of RNA (Feng, Li et al., 2020, Hennig, Kong et al., 2015, Hentze, Castello et al., 2018, Knott, Bond et al., 2016). NONO is a member of the highly conserved Drosophila behaviour/human splicing (DBHS) protein family, which also includes the splicing factor proline/glutamine rich (SFPQ) and paraspeckle protein component 1 (PSPC1) proteins. DBHS proteins have two N-terminal RNA recognition motifs (RRM), a NonA/paraspeckle domain (NOPS), a coiled-coil at the C-terminus and N- and C-terminal intrinsically disordered regions (IDRs). DBHS proteins form obligate dimers that can bind DNA, RNA, undergo oligomerisation, mediate additional protein-protein interactions and also undergo liquid-liquid phase separation. Combined, these different interactions suggest DBHS proteins act as ‘molecular scaffolds’ to carry out their multipurpose activities in many facets of gene regulation.

Several studies have profiled the genome, and transcriptome wide DNA and RNA substrates bound by NONO in different biological contexts (Benegiamo, Mure et al., 2018, Ma, Karwacki-Neisius et al., 2016, Van Nostrand, Freese et al., 2020, Xiao, Chen et al., 2019). Broadly these studies have revealed widespread binding to diverse gene regulatory elements in chromatin as well as binding to mainly intronic elements of pre-mRNAs. In some instances this binding is linked to specific co-regulatory networks such as NONO binding to ERK promoter targets in stem cells (Ma et al., 2016), or binding to pre-mRNA of transcripts coding metabolic genes in liver hepatocytes (Benegiamo et al., 2018). Whilst NONO/ERK association at chromatin is required for mouse embryonic stem cell (mESC) pluripotency, there is little mechanistic insight explaining the consequences of NONO binding to RNA in different contexts. Other DBHS proteins have also been analysed by CLIP and ChIP (Hosokawa, Takeuchi et al., 2019, Iida, Hagiwara et al., 2020, Stagsted, O’Leary et al., 2021, Takeuchi, Iida et al., 2018); for instance, SFPQ binds to and enables processing of long introns in neurons, and prevents intron retention in ALS motor neurons (Luisier, Tyzack et al., 2018).

Given their extensive and diverse DNA and RNA targets, DBHS proteins are found throughout the nucleus as well as in specific sub-nuclear sites, such as paraspeckles – where they bind and play an essential role stabilising the lncRNA NEAT1_2 scaffold (Knott et al., 2016) – and to sites of DNA damage (Krietsch, Caron et al., 2012, Li, Li et al., 2014). Both paraspeckles and DNA damage foci are now classed as condensates built by the liquid-liquid phase separation properties of various component proteins (Fox, Nakagawa et al., 2018, Spegg & Altmeyer, 2021). Liquid-liquid phase separation is an emerging phenomenon explaining the dynamic association of molecules, including RNA binding proteins, into membrane-less organelles, or condensates (Alberti & Dormann, 2019, Sabari, Dall’Agnese et al., 2018, Zbinden, Pérez-Berlanga et al., 2020). Recently it was demonstrated that NONO undergoes phase separation at DNA damage foci (Fan, Wang et al., 2021), however how phase separation by NONO plays a role in gene regulation is unknown.

Super-enhancer (SE) regulated gene networks are defined by ChIP signatures and establish distinct cell lineage programs. Recently it was revealed that super enhancers are controlled by the formation of phase separated condensates composed of transcription factors and transcriptional cofactors (Boija, Klein et al., 2018, Guo, Manteiga et al., 2019). This regulation may be critical for neuroblastoma, as two main SE-associated transcriptional networks control lineage identity, intra-tumoral heterogeneity and cell-type specific gene expression, including a mesenchymal cell state and a noradrenergic cell state (Boeva, Louis-Brennetot et al., 2017, van Groningen, Koster et al., 2017). The noradrenergic cells are further subdivided into three major SE-driven epigenetic subtypes and their underlying master regulatory networks recapitulating three clinical groups in neuroblastoma tumours and cell lines (Gartlgruber, Sharma et al., 2021). A small number of key transcription factors associated with SE were identified as members of the transcriptional core regulatory circuitry (CRC) that determine noradrenergic cell fate and growth, such as MYCN, PHOX2B, HAND2, GATA2 and GATA3 (Durbin, Zimmerman et al., 2018, Gartlgruber et al., 2021, van Groningen et al., 2017).

In this study, we take a holistic look at NONO in neuroblastoma to determine the mechanisms linking NONO to poor outcome in this context. We combine cell imaging, biophysical analysis, and various RNA-seq analyses to reveal a complex picture of NONO-dependent regulation of gene expression. We find NONO in numerous small non-paraspeckle condensates throughout the nucleus, tethered to these by RNA and DNA. Accordingly, NONO mutants that cannot bind RNA mis-localise in larger spherical non-functional puncta that more readily phase separate, confirming the critical role of NONO RNA binding in its function. Using PAR-CLIP we show NONO binds to the 5’ ends of pre-mRNA and influences pre-mRNA processing. Notably, NONO-bound transcripts are also more likely to be SE-regulated, including HAND2 and GATA2. We find that the decreased expression of HAND2 and GATA2 after NONO depletion is likely mediated by inefficient pre-mRNA processing at these loci. Thus overall, NONO requires the coordinated integration of multilevel components of mechanistic processes and signals to enact its oncogenic program.

## Results

### NONO puncta are condensates dependent on RNA and DNA

Given that high levels of NONO are correlated with poor patient outcome in neuroblastoma (Fig 1A and (Liu et al., 2014)), we set out to investigate the role of NONO in this biological context, to inform future rational design of therapeutics. At the cellular level, we have previously demonstrated that paraspeckles – as illustrated by Fluorescence In Situ Hybridisation (FISH) against NEAT1_2 – are co-localized with a subset of NONO immunofluorescence signal (Naveed, Cooper et al., 2021, Yamazaki, Souquere et al., 2018). When comparing the distribution of NONO puncta between high-risk neuroblastoma (KELLY and BE(2)-C) and HeLa cell lines, we observed fewer paraspeckles in neuroblastoma cells (Fig 1B), as shown previously (Naveed et al., 2021). Instead of within paraspeckles, NONO is localized in numerous small puncta throughout the nucleus, in both neuroblastoma cell lines.

**Figure 1:**
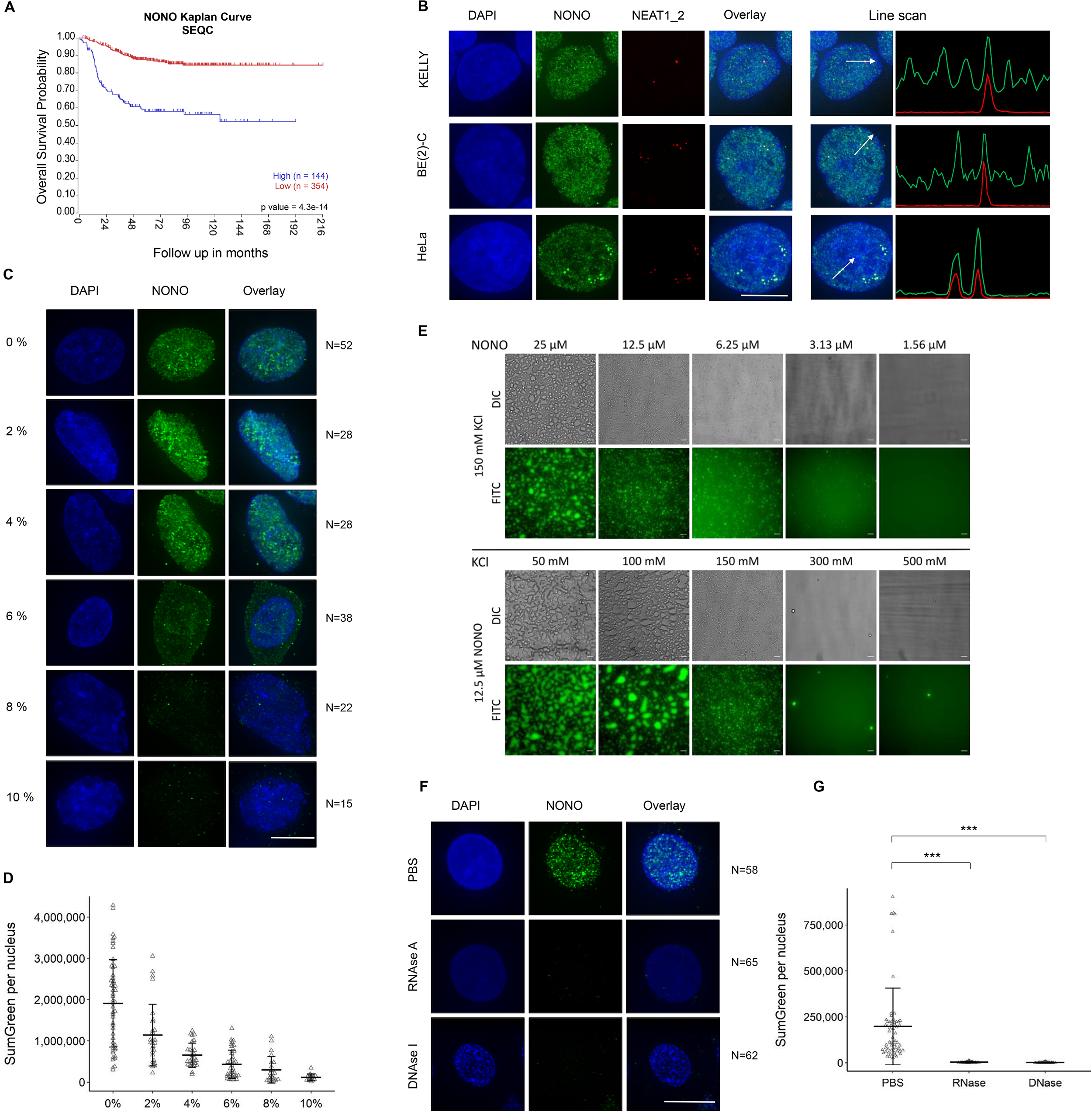
Both RNA and DNA are essential for distinct distribution of NONO puncta in neuroblastoma cell lines. (A) The probability of overall survival is lower in neuroblastoma patients with high NONO expression based on Kaplan-Meier curve using the SEQC neuroblastoma dataset. (B) Fluorescence micrograph images of representative cells stained for NONO and NEAT1_2 in KELLY and BE(2)-C neuroblastoma and Hela cells showing clear paraspeckle (as marked by NEAT1_2), and non-paraspeckle NONO puncta. DAPI (blue) stain indicates cell nuclei, NONO immunofluorescence (green) and NEAT1_2 RNA FISH (red). Scale bar: 5 μm. (C) Fluorescence micrograph images of representative cells stained for NONO in KELLY cells treated with 2, 4, 6, 8 or 10% 1,6 hexanediol showing dissolution of NONO puncta with increasing concentration. Scale bar: 5 μm. (D) Dot plot of summed green fluorescence per nucleus at different concentrations of 1,6 hexanediol as in (C). Bars are SD. (E) Recombinant GFP-NONO_WT can phase separate spontaneously. Its propensity to phase separate increases with increasing protein and decreasing KCl concentration. Scale bar: 20 μm. (F) Fluorescence micrograph images of representative cells stained for NONO in KELLY cells treated with PBS, RNase A or Dnase I, as indicated. Scale bar: 5 μm. (G) Dot plot of summed green fluorescence per nucleus in (F). Bars are SD. ***P<0.001.

To address the nature of the small non-paraspeckle NONO puncta, we incubated cells with 1,6-hexanediol (a compound that disrupts liquid-liquid phase separated condensates) and observed dramatically reduced NONO signal intensity in KELLY (Fig 1C-1D) and HeLa (Fig S1A-S1B) cells. Supporting a role for phase separation in the formation of NONO puncta, we also observed that recombinant full-length GFP tagged NONO could form droplets *in vitro*, regulated by varying concentrations of either NONO protein or KCl (Fig 1E). In addition, nuclease digestion using either RNase A or DNase I completely eradicated the nuclear NONO puncta signal in KELLY (Fig 1F-1G) and HeLa (Fig S1C-S1D) cells. Together, these *in vitro* and *in vivo* observations suggest that the numerous non-paraspeckle NONO puncta may be phase-separated condensates that are dependent on both RNA (distinct from NEAT1_2) and DNA for their structural integrity.

### RNA recognition motif 1 (RRM1) is essential for NONO to bind RNA targets

Given the importance of RNA in NONO condensate formation, we next addressed NONO RNA binding ability for its localisation and function. The canonical RRM1 is structurally characterized in NONO, and is required for NONO binding RNA *in vitro*, however its role in RNA binding has not been fully evaluated in different biological settings (Fox, Bond et al., 2005, Knott et al., 2016, Knott, Chong et al., 2021, Kuwahara, Ikei et al., 2006, Passon, Lee et al., 2012). Thus, we compared the localisation of overexpressed YFP fused wild-type NONO (YFP-NONO_WT) with NONO lacking the RRM1 (YFP-NONO_ΔRRM1). We observed that, compared to YFP-NONO_WT, YFP-NONO_ΔRRM1 no longer co-localized with paraspeckles in KELLY (Fig 2A-2B) and HeLa (Fig S2A-S2E) cells suggesting the loss of RRM1 abrogated the ability to bind NEAT1_2. The YFP-NONO_ΔRRM1 puncta were also more spherical, fewer in number, and larger, compared to YFP-NONO_WT (Fig 2C-2E). Functionally, intact RRM1 in NONO was linked to KELLY cell proliferation, as measured by the percentage of BrdU positive cells increasing when exogenous YFP-NONO_WT was overexpressed, but not YFP-NONO_ΔRRM1 (Fig 2F). We also mutated some additional residues in the NONO NOPS domain that may mediate RNA association, as identified in RBDMap RNA crosslinking studies (Castello, Fischer et al., 2016). However, whilst these point mutants displayed a similar localisation to YFP-NONO_ΔRRM1, this is more likely the result of failure to dimerize than bind RNA, as, unlike YFP-NONO_WT, or YFP-NONO_ΔRRM1, the YFP-NONO point mutants no-longer co-immunoprecipitated endogenous SFPQ (Fig S2F).

**Figure 2:**
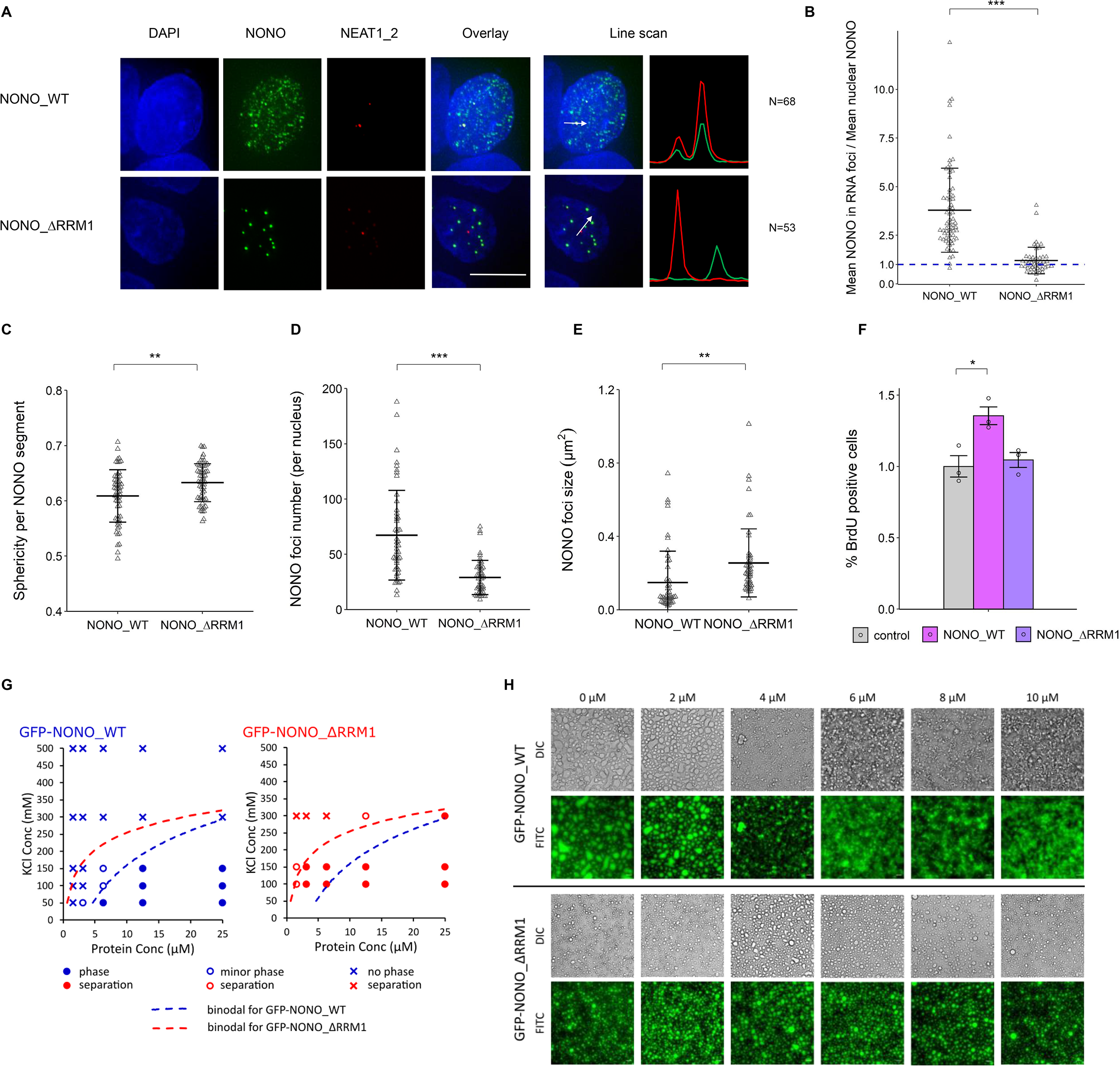
RRM1 is an important region for NONO to bind RNA targets in KELLY cells. (A) Fluorescence micrograph images of representative cells stained for NONO and NEAT1_2 after transfection with plasmids expressing YFP fused NONO_WT and NONO_ΔRRM1 protein. DAPI (blue) stain indicates cell nuclei, YFP fused NONO (green) and NEAT1_2 RNA FISH (red). Scale bar: 5μm. (B) The enrichment of mean NONO fluorescence detected within RNA FISH foci is quantitatively determined as a ratio relative to mean nuclear NONO fluorescence in (A). Bars are SD. *p<0.05, ** p<0.01, ***p<0.001. (C) Sphericity per NONO segment, (D) NONO foci number per nucleus and (E) NONO foci size between YFP fused NONO_WT and NONO_ΔRRM1 plasmids were measured. Bars are SD. (F) Percentage of BrdU incorporation between control cells (YFP only transfection), cells with YFP fused NONO_WT and YFP-NONO_ΔRRM1 plasmids transfected. Bars are SEM. n≥3. (G) Recombinant GFP-NONO_ΔRRM1 has a higher propensity to phase separate than GFP-NONO_WT at the same KCl concentration. Blue and red colours denote GFP-NONO_WT and GFP-NONO-ΔRRM1 respectively. Filled circles, open circles and crosses indicate distinct phase separation, minor phase separation and no phase separation respectively. Dotted lines denote putative binodal line separating single and two phase states. Scale bar: 20 μm. (H) PS-ASO against NEAT1 disrupts the phase separation of GFP-NONO_WT but not GFP-NONO_ΔRRM1 in a PS-ASO concentration-dependent manner. Scale bar: 20 μm.

*In vitro*, recombinant GFP-NONO_ΔRRM1 could form droplets, with an increased propensity for droplet formation compared to GFP-NONO_WT (Fig 2G). Moreover, adding 2’-O-methyl phosphorothioate antisense oligonucleotides (PS-ASO) against NEAT1 (a known RNA substrate for NONO (Vickers, Rahdar et al., 2019)) into GFP-NONO_WT solution caused NONO droplets to become small fibrils (Fig 2H, top). In contrast, adding PS-ASO to GFP-NONO_ΔRRM1 had no effect on droplets, indicating NONO_ΔRRM1 is impervious to the addition of nucleic acid NONO substrate (Fig 2H, bottom). Taken together, these results suggest RNA binding through RRM1 modulates the propensity for NONO to phase separate, altering sub-nuclear distribution.

### NONO binds predominantly within introns

We next assessed the transcriptome-wide binding of NONO in KELLY and BE(2)-C neuroblastoma cells. We performed PAR-CLIP and found, similar to other DBHS protein CLIP experiments, the majority of NONO binding occurs within introns, even when normalizing for the percentage of the transcriptome made up of introns (Fig 3A). NONO binding was biased towards the 5’ end of transcripts (Fig 3B), a pattern also previously observed in CLIP against NONO and other DBHS family proteins in other biological settings (Benegiamo et al., 2018, Jiang, Shao et al., 2017, Takayama, Suzuki et al., 2017). To further define NONO targets, we summed aligned reads containing T-to-C transitions across genes (Hafner, Landthaler et al., 2010). Using TPM (transcripts-per-million (Wagner, Kin et al., 2012)) and normalizing for gene length, NONO targets (the top 20 percentile by TPM of all genes containing T-to-C read alignments, 1831 genes) were enriched in several ontologies, including ‘mRNA binding’ and ‘post-transcriptional gene silencing’ in KELLY and BE(2)-C cells (Fig 3C-3D).

**Figure 3:**
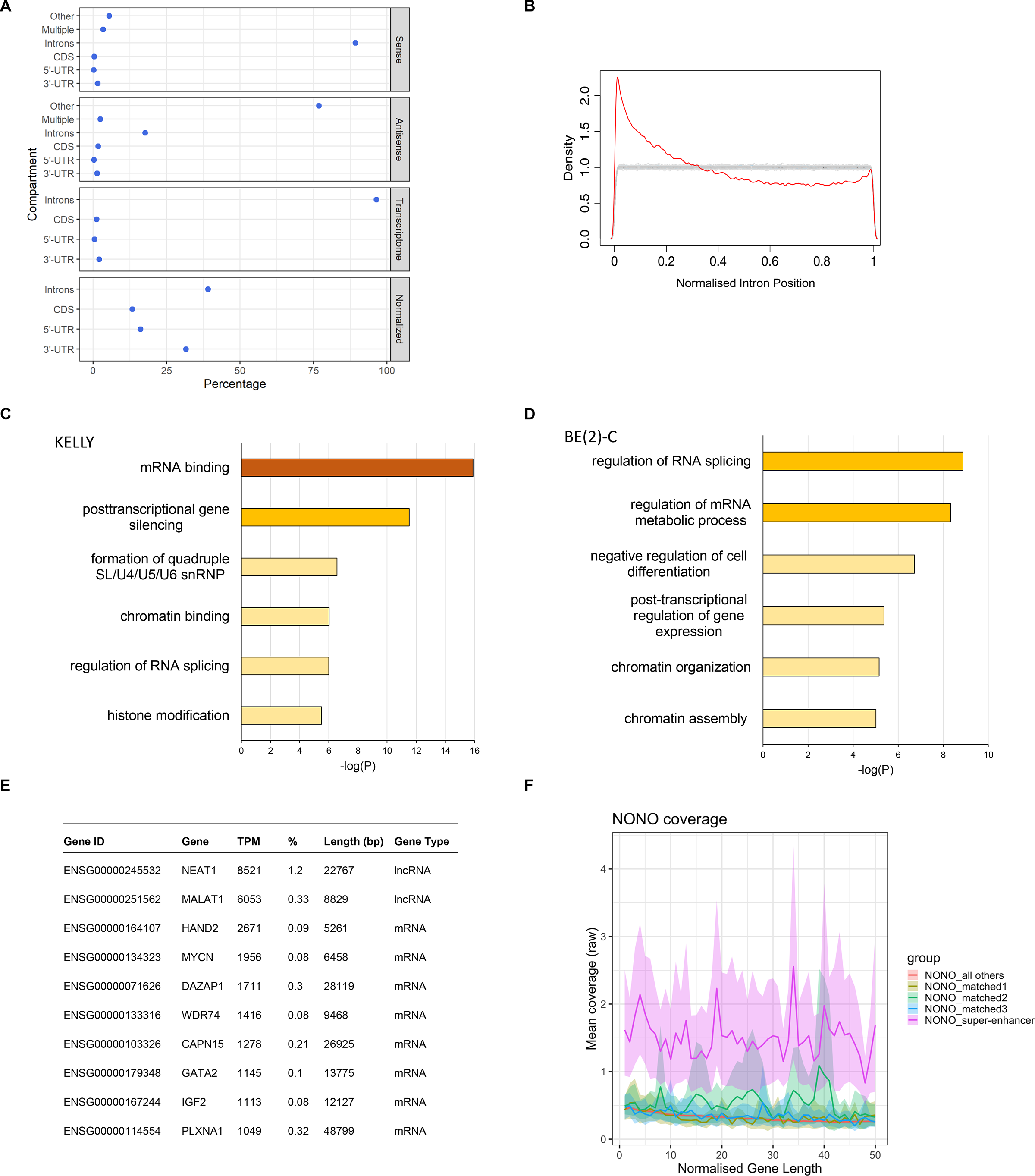
NONO preferentially binds 5’ introns and SE regulated target genes have more NONO binding. (A) NONO RNA binding sites are strongly biased towards introns, as determined by PAR-CLIP and wavClusteR. Sense: percentage of annotated binding sites on sense strand. Antisense: percentage of annotated binding sites on antisense strand. Transcriptome: relative length of each annotation category within the transcriptome. Normalized: percentage of annotated binding sites corrected by total length of each annotation category in transcriptome. (B) NONO binding sites within introns show preference for the 5’ end of genes as determined by PAR-CLIP and PARpipe. (C) Metascape gene ontology analysis of NONO-bound target transcripts. (D) Metascape gene ontology analysis of NONO-bound target transcripts in BE(2)-C cells. (E) Summary of most highly NONO-bound RNA targets by NONO TPM and % of all T-to-C reads. (F) NONO PAR-CLIP coverage profiles (metagene2) across SE-regulated genes compared to expression and length-matched controls. NONO binds to SE regulated target genes with greater coverage compared to matched controls.

Despite NEAT1 being less abundant in KELLY cells than other cell types, it was nevertheless the top NONO target RNA (Fig 3E), with by far the largest percentage of PAR-CLIP reads of any RNA, reflecting the high affinity of NONO for NEAT1 (Naveed et al., 2021). Amongst the rest of the top 10 transcripts, HAND2 and GATA2 were notable as they encode super-enhancer (SE) regulated transcription factors essential to mediate lineage control in neuroblastoma (Boeva et al., 2017, van Groningen et al., 2017). To determine if other SE-regulated genes were also highly bound by NONO, we analyzed publicly available H3K27 acetylation ChIP-seq data from Chipumuro *et al*. (Chipumuro, Marco et al., 2014) to identify the SE profile in KELLY cells, examining the overlap of NONO RNA binding for a subset of genes within SE regions from the H3K27ac ChIP-seq analysis. We found that transcripts from genes within SE regions had substantially greater NONO RNA binding when compared with expression-matched controls, suggesting a preferential RNA binding of NONO to SE-regulated target gene transcripts (Fig 3F).

To validate our PAR-CLIP findings, we conducted RNA immunoprecipitation (RIP) against NONO followed by RNA quantification of mRNAs and pre-mRNAs for MYCN, DAZAP1, GATA2 and HAND2, observing greater NONO binding to pre-mRNAs (Fig 4A). We then used siRNA to knock down endogenous NONO and rescued with siRNA-resistant YFP-NONO_WT, or YFP-NONO_ΔRRM1 plasmids. When compared to controls, various target transcripts including total_NEAT1, DAZAP1, pre_HAND2 and pre_KCNQ2 were enriched with exogenous YFP-NONO_WT, but not with YFP-NONO_ΔRRM1, confirming that binding to pre-mRNA targets is lost in the mutant NONO (Fig 4B). The pre-HAND2 and pre_KCNQ2 transcripts were also significantly different between the cells with YFP-NONO_WT and YFP-NONO_ΔRRM1. To determine if any NONO condensates corresponded to these target transcripts we used RNA FISH probes against NEAT1, or pre-mRNAs of GATA2 and HAND2, in conjunction with immunofluorescence against NONO. Whilst each target transcript only had 1-2 foci per cell, likely representing nascent transcripts forming near gene loci, nevertheless, these foci were highly enriched in NONO (Fig 4C-4D). Thus, nuclear NONO condensates represent sites of NONO binding to a variety of lncRNA and pre-mRNA targets, particularly within the 5’ part of pre-mRNAs regulated by super enhancers.

**Figure 4:**
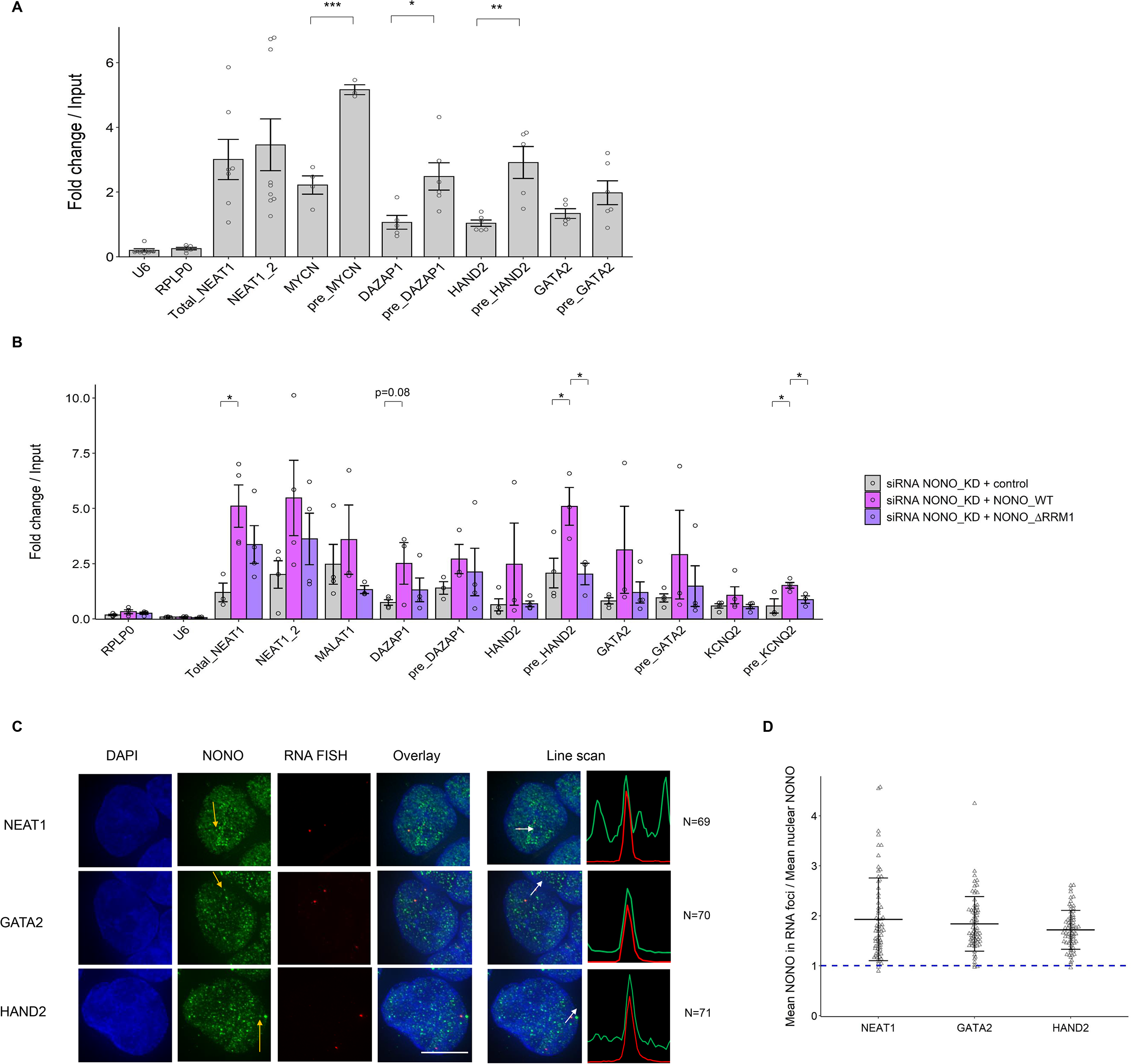
NONO binds more abundantly on pre-mRNAs than mature counterparts in KELLY cells. (A) Relative pre-mRNA levels and their mature counterparts were measured by NONO RNA RIP followed by RT-qPCR. RIP data with normal mouse serum IgG as controls are not displayed because all values are <0.01. Bars are SEM. n≥3. *p<0.05, ** p<0.01, ***p<0.001. Relative pre-mRNA levels and their mature counterparts were measured by NONO RNA RIP followed by RT-qPCR in cells transfected sequentially with NONO KD siRNA and then siRNA-resistant control (YFP only), YFP fused NONO_WT or NONO_ΔRRM1 plasmids. Bars are SEM. n≥3. (C) Fluorescence micrograph images of representative cells stained for NONO and NEAT1_2 (top), GATA2 (middle) and HAND2 (bottom). DAPI (blue) stain indicates cell nuclei, NONO immunofluorescence (green) and RNA FISH (red) for NEAT1_2, GATA2 and HAND2. Scale bar: 5μm. (D) In micrograph image quantitation analysis, the enrichment of mean NONO fluorescence detected within RNA FISH foci is determined as a ratio relative to mean nuclear NONO fluorescence in (C). Bars are SD.

### NONO maintains proper RNA processing and splicing at the 5’ end of transcripts

To address the consequence of NONO knockdown (KD) on target gene expression, we used two siRNAs against NONO followed by RNA-seq in KELLY cells. We first confirmed a significant reduction in NONO mRNA and protein levels after KD (Fig 5A) and then compared four controls with eight NONO KD samples (4 for each NONO siRNA) using RNA-seq. In total, 3594 genes were differentially expressed (DESeq2 p_adj_ < 0.1), with no bias for up- or down-regulation (Fig 5B) and no obvious difference according to gene length (Fig S3A). Genes upregulated following NONO KD were enriched for ontologies relating broadly to the negative control of transcription, whereas genes downregulated were enriched in pathways relating to cholesterol synthesis and metabolism (Fig S3B-S3C). Gene Set Enrichment Analysis (GSEA) also found the most highly enriched pathways among NONO KD samples (containing genes that increased in activity) were GO:0006342 ‘chromatin silencing’ (GSEA p<10^-3^, FDR<10^-3^) and GO:0045814 ‘negative regulation of gene expression (epigenetic)’ (GSEA p<10^-3^, FDR<10^-3^).

**Figure 5:**
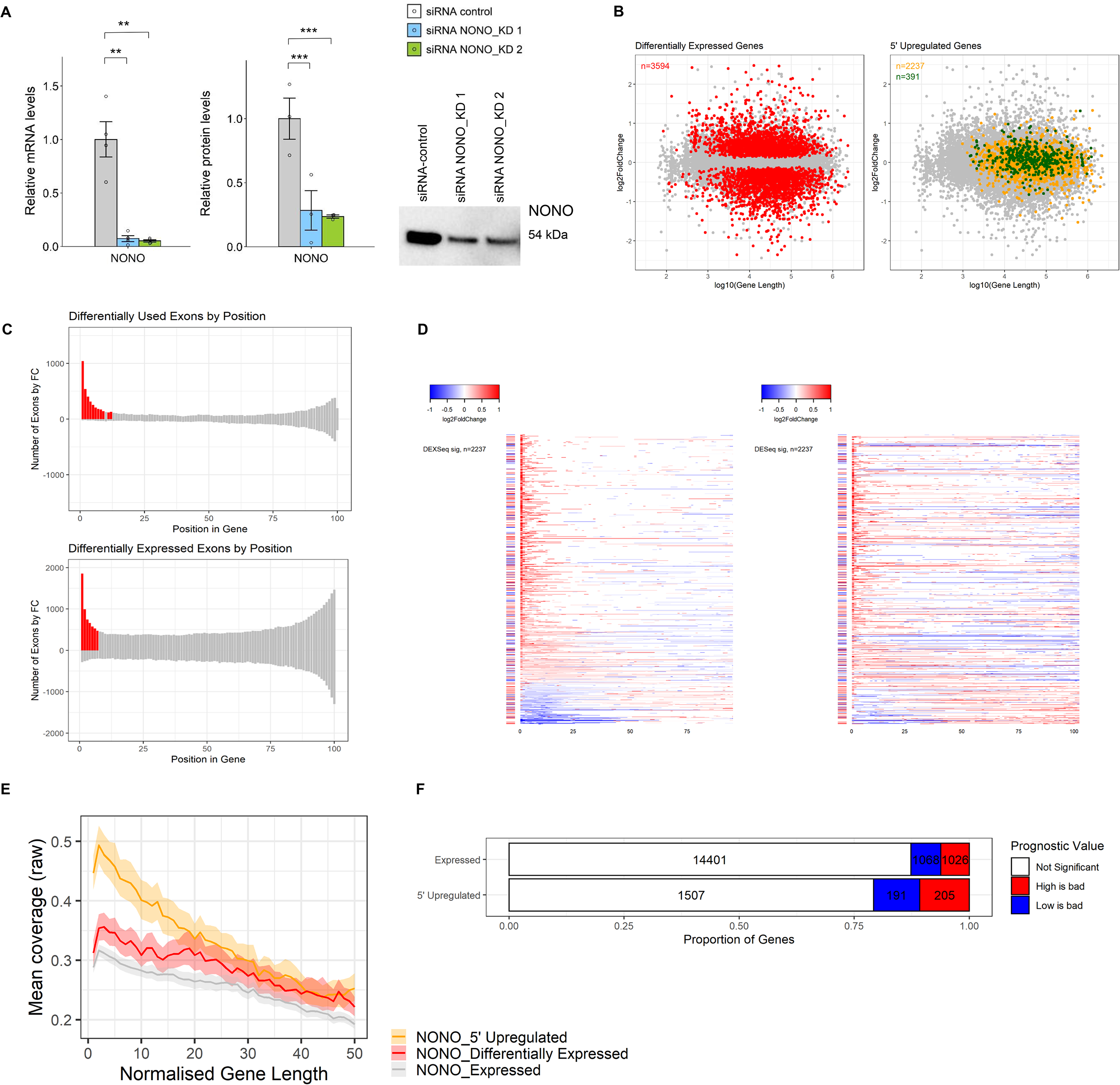
NONO KD results in altered RNA processing and splicing in KELLY cells. (A) Relative NONO mRNA and protein levels in KELLY cells treated with control or NONO KD siRNAs. Representative Western blot image for NONO protein. Bars are SEM. n≥3. ** p<0.01, ***p<0.001. (B) Differentially expressed genes do not demonstrate a length, or fold-change bias (red). The upregulated genes at the 5’ end (orange), including those that are NONO bound (green), have a moderate bias towards longer genes. (C) Histogram of significant differential usage and differential expression events by gene position. The proportion of total significant events for each exon bin is compared to the median proportion. Bins whose proportion exceeds the median by twice the standard deviation are coloured red. (D) A subset of genes show 5’ upregulation of both differential usage (left, DEXSeq) and differential expression (right, DESeq2). Each row represents a gene, split into 100 bins (exons and introns). Coloured bins represent significant events in differential usage and differential expression testing. The bar to the left of each row indicates whether the gene as a whole is differentially expressed. Red indicates increased expression, and blue indicates decreased expression. (E) NONO coverage across genes showing 5’ upregulation has a distinct 5’ bias when compared with genes which are differentially expressed, and compared to all expressed genes (metagene2). (F) There is a higher proportion of neuroblastoma-prognostic genes in the upregulated genes at the 5’ end compared with the other expressed genes based on the Kaplan-Meier tool using four neuroblastoma datasets (Cangelossi, Maris, SEQC and Versteeg).

Given NONO’s significant RNA binding activity, we next assessed the role of NONO in alternative splicing. To identify any overall changes we used IsoformSwitchAnalyzR, which showed that NONO disruption induced a significant bias in the use of upstream ‘alternative transcription start sites’ (ATSS) and downstream ‘alternative 3’ end acceptor sites’ (A3) (Fig S3D). To further focus on these splicing changes, we used DESeq2 to test changes in the absolute expression of each intron and exon between control and KD samples. In addition, we used DEXSeq to test changes in the expression of each intron and exon relative to the overall expression of the parent gene: ‘differential usage’. Thus, DEXSeq tests for changes that occur within a gene, controlling for expression changes of the gene as a whole. To look for a pattern of regulatory change with NONO KD, we divided each gene into 100 bins and looked at the position of each differentially expressed and differentially used exon/intron. We tested whether the proportion of significant upregulation events was significantly greater than the median, finding that exons within the first 13 bins had a significantly greater proportion of upregulated usage events (Fig 5C, top) and exons within the first 7 bins had a greater proportion of significantly upregulated expression changes (Fig 5C, bottom).

Examining the full transcripts (exons and introns) of the 2237 genes which had significant expression and/or usage events identified in Fig 5C, there is a significant bias for 5’ upregulated usage and expression (Fig 5D, red). However, despite this 5’ difference, there is no consistent change in the overall transcript expression for these genes (Fig 5B, yellow, Fig 5D row-side colours), even when overlapping with top NONO hits (Fig 5B, green). Thus, we observed that NONO KD induces upregulated usage and expression at the 5’-most extent of genes, which is independent of overall expression changes. This suggests a deficiency in processing at the 5’ end of transcripts in the absence of NONO.

Combining the NONO KD RNA-seq with NONO PAR-CLIP, NONO-bound RNAs were significantly more likely to be differentially expressed (χ^2^ = 76.9, p < 10^-15^, Fig S3E). To test whether NONO RNA binding is implicated in the observed 5’ usage and expression changes, we looked at the average NONO PAR-CLIP coverage across these genes. The selected genes displayed a pronounced binding bias at the 5’ end compared to genes which were differentially expressed overall, yet showed approximately equivalent binding at the 3’ end (Fig 5E). Based on survival analysis, we also found that the upregulated genes at the 5’ end are more likely to be prognostic in neuroblastoma, including GATA2 and HAND2, compared with other expressed genes (Fig 5F). These results suggest that NONO RNA binding at the 5’ end may be required for successful pre-mRNA processing of transcripts with important roles in neuroblastoma.

### Delayed RNA processing of GATA2 and HAND2 results in decreased expression

Given that GATA2 and HAND2 have been demonstrated to modulate differentiation and migration in neuroblastoma (Voth, Oberthuer et al., 2009, Willett & Greene, 2011), we examined if their expression levels were regulated by NONO in KELLY cells. Indeed, GATA2 and HAND2 mRNA and protein levels were decreased after NONO KD (Fig 6A-6C).

**Figure 6:**
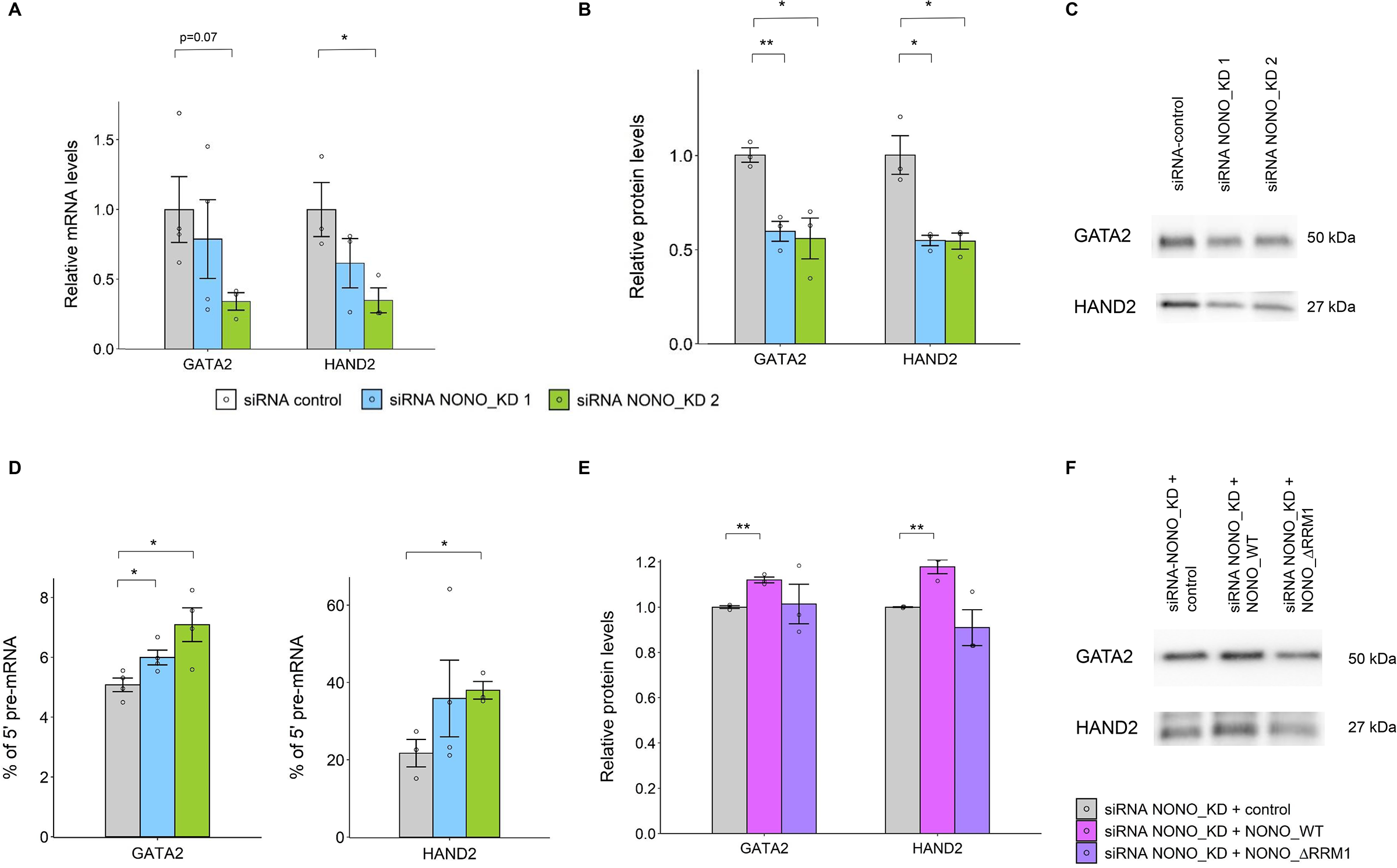
NONO KD reduces mRNA and protein expression of GATA2 and HAND2, which may result from inappropriate processing and splicing in KELLY cells. (A) Relative GATA2 and HAND2 mRNA levels via RT-qPCR between control and NONO KD siRNAs. Bars are SEM. n≥3. *p<0.05, ** p<0.01. (B) Western blot quantitation analysis of GATA2 and HAND2 protein levels between control and NONO KD siRNAs. Bars are SEM. n≥3. (C) Representative Western blot images for GATA2 and HAND2 proteins in (B). (D) The percentage of GATA2 and HAND2 pre-mRNAs at the 5’ end was measured relative to their mature transcripts between control and NONO KD siRNAs. Bars are SEM. n≥3. (E) Western blot quantitation analysis of GATA2 and HAND2 protein levels in cells transfected sequentially with NONO KD siRNA and then siRNA-resistant control (YFP only), YFP fused NONO_WT or NONO_ΔRRM1 plasmids. Bars are SEM. n≥3. (F) Representative Western blot images for GATA2 and HAND2 proteins in (E).

We next determined the possible mechanisms underlying the decreased expression of GATA2 and HAND2. Although we could detect NONO binding to the *GATA2* gene by ChIP-qPCR, this chromatin binding did not alter when GATA2 transcription was reduced (Fig S4A-S4C), suggesting the binding of NONO to DNA and nascent RNA were not linked. We also tested any NONO-dependency for transcriptional elongation of GATA2 and HAND2, by ChIP-qPCR of phosphorylated Serine 2 of RNA polymerase II, yet found no difference in signal with NONO KD (Fig S4D), and no co-localisation of NONO condensates with PolII-phospho-Ser2 condensates by immunofluorescence (Fig S4E-S4F). In contrast, we found that NONO KD induced a higher relative expression of GATA2 and HAND2 pre-mRNAs at the 5’ end when compared to their mature transcripts (Fig 6D). Furthermore, overexpression of NONO_WT, rather than NONO_ΔRRM1 led to increased GATA2 and HAND2 protein levels (Fig 6E-6F), indicating that NONO has an effect on target gene expression through adequate RNA binding activity. Taken together, these data indicate that NONO binds pre-mRNAs and enhances RNA processing and splicing close to the 5’ end of transcripts, driving optimal oncogenic expression levels.

## Discussion

NONO acts as a molecular scaffold in gene regulation in many contexts. In this study, we identified NONO-mediated enhancement of RNA processing at the 5’ end of important transcripts as a key molecular mechanism in neuroblastoma. This 5’ associated RNA processing activity is linked to NONO nuclear condensates that form at individual gene loci, including those of the super-enhancer regulated genes GATA2 and HAND2. In the absence of functional NONO-RNA condensates, GATA2 and HAND2 protein levels decrease, with evidence for stalled 5’ RNA processing. Therefore, we propose a model whereby NONO binds to, and coats the 5’ ends of nascent transcripts, forming gene-body splicing-associated condensates to enhance gene expression and support an oncogenic program.

Here we show NONO forms numerous non-paraspeckle phase separated condensates in the nucleus, and that NONO readily forms droplets *in vitro*. NONO contains IDRs at the N- and C- termini, with a central globular domain for RNA binding, dimerization and coiled-coil oligomerisation; however, it is not known which of the IDRs, if any, drive condensate formation (Knott et al., 2016, Knott, Lee et al., 2015, Passon et al., 2012). Recently it was shown that the DBHS protein PSPC1 requires the C-terminal IDR to undergo phase separation (Li, Cui et al., 2021), and the analogous IDR exists in NONO, therefore this would be a good candidate region to test. We also showed that RNA binding, via RRM1, attenuates NONO phase separation, such that mutants with impaired RNA binding ability more readily phase separate *in vitro* and form large spherical droplets in the nuclei of cells. In contrast, wildtype NONO binds RNA and forms small fibrils *in vitro* and much smaller, finer, irregular condensates inside the nucleus. The propensity of RNA to prevent aberrant, gross phase separation by NONO is similar to what is observed for FUS, where mutants lacking RNA binding capacity excessively phase separate, while the wild-type protein is mostly diffuse (Maharana, Wang et al., 2018). However, NONO is different to FUS in that wildtype NONO is not diffuse, but instead forms many hundreds of smaller condensates, each likely representing a site of nascent transcription. For FUS, addition of small amounts of RNA promotes phase separation into droplets *in vitro*, whereas high RNA levels prevent droplet formation (Maharana et al., 2018). In contrast, addition of increasing concentration of RNA *in vitro* causes NONO droplets to convert into small fibrils, a behaviour also observed for PSPC1 (Shao, Bi et al., 2022). Thus, distinct families of RNA binding proteins use their suite of multivalent interactions to respond differently to the presence of RNA *in vitro* and *in vivo*. Our work adds to the growing appreciation of how RNA binding domains influence the final material state and modulate the dynamics of condensates in general (Gotor, Armaos et al., 2020, Wiedner & Giudice, 2021).

Paraspeckles are well-known protein-RNA condensates that form through microphase separation and are also not typically spherical. In the two main steps of paraspeckle formation, first, DBHS proteins, including NONO, bind and stabilise NEAT1 lncRNA. Secondly, FUS binds and carries out phase separation dependent on its prion-like domain (Yamazaki, Yamamoto et al., 2021). Whilst a role for NONO-driven phase separation in paraspeckle formation is yet to be established, NONO phase separation is required for radiation-induced DNA damage repair (Fan et al., 2021). In addition, NONO also promotes phase separation and activation of TAZ, the hippo pathway effector, to drive oncogenic transcription and tumorigenesis in glioblastoma, although a direct role for phase separation of NONO itself was not addressed in that study (Wei, Luo et al., 2021). Thus, NONO phase separation may be important for many of its regulatory activities in different contexts.

As well as binding to RNA, there is evidence from us and others, that NONO is bound to chromatin, albeit not through direct binding in many cases (Knott et al., 2016, Yang, Yang et al., 1997). In development, NONO and SPFQ are enriched at bivalent genes with high levels of poised RNA PolII, influencing lineage commitment (Ma et al., 2016, Van Nostrand et al., 2020, Xiao et al., 2019, Yadav, Hao et al., 2014). NONO is also responsible for recruitment of the 5-hydroxymethylcytosine enzyme TET1 to chromatin in mESCs (Li, Karwacki-Neisius et al., 2020). NONO RRM1 deletion did not abrogate its interaction with TET1, nor prevent NONO-dependent recruitment of TET1 to chromatin in mESCs, therefore NONO RNA binding appears to be less important in regulation of pluripotency. This is in contrast to PSPC1, which recruits TET2 to chromatin, dependent on its RNA binding activity (Guallar, Bi et al., 2018). However, for neuroblastoma our data argue that chromatin associated NONO is not the main driver of the gene regulatory role in this context, but instead might have a permissive role, preparing and facilitating a rapid transcriptional response when stimulators are present. This interpretation is partly due to our ChIP evidence that NONO recruitment to the GATA2 promoter is insensitive to transcription inhibition. Further, NONO mutants lacking RNA binding were no longer recruited to normal NONO condensates, instead appearing in fewer, larger, rounder nuclear droplets that are non-functional as they do not support enhanced cell proliferation in a similar manner to wildtype NONO.

Our sequencing and CLIP data suggest many genes display NONO-dependent expression. These genes normally have high levels of NONO bound at the 5’ ends of their transcripts, principally in the introns. In the absence of NONO there is an increase in reads corresponding to the first ∼13% of transcript length, however this does not always correlate with overall differential expression. In the case of GATA2 and HAND2 we showed NONO loss leads to decreased protein production, with an increase in 5’ end of transcripts corresponding to the pre-mRNA, implicating improper RNA processing in this downregulation. Thus, we propose a model in which a gene regulatory role for NONO in neuroblastoma hinges on binding to the pre-mRNA of nascent SE-regulated genes, promoting the formation of RNA-processing condensates, allowing efficient splicing. Extensive prior literature supports a role for NONO in pre-mRNA processing and splicing in different cell types and clinical samples (Feng et al., 2020, Yamamoto, Osawa et al., 2018). Although NONO is not a crucial component in spliceosome assembly, it interacts with critical spliceosomal proteins (Kameoka, Duque et al., 2004, Zhang & Wu, 1996).

The primary mechanism of RNA processing enhanced by NONO condensates is still uncertain. Preventing intron retention is one explanation. However, using established bioinformatic pipelines we did not observe intron retention to be significantly altered with NONO KD. Given the 5’ usage bias, this may be due to an inability to differentiate first intron retention, from all-intron retention. Others have shown NONO KD resulted in aberrant splicing, including intron retention, and altered gene expression in mouse developing retina (Yadav et al., 2014). Moreover, SFPQ, but not NONO, prevents intron retention in early neural differentiation (Luisier et al., 2018, Stagsted et al., 2021). Another factor to consider is the importance of NONO condensates for intron removal in co-transcriptional splicing, as opposed to post-transcriptional splicing. Whilst earlier findings support a role for NONO in co-transcriptional splicing (through association both with nascent RNA and PolII CTD) (Emili, Shales et al., 2002, Kameoka et al., 2004), new evidence of the importance of ‘nuclear anchoring’ of partially processed, but fully transcribed, pre-mRNA transcripts at the gene locus is emerging (Girard, Will et al., 2012, Popp & Maquat, 2013, Quinn & Chang, 2016). In this context, a chromatin-anchored nuclear pool of partially spliced, but polyadenylated RNA, may act in a regulatory manner as a reservoir for mature mRNA, upon splicing. Intriguingly, such a mechanism seems to be important in the neuronal gene regulation context (Yeom, Pan et al., 2021). Important future experiments would therefore include testing if NONO condensates act at the co- or post-transcriptional level, by repeating NONO KD RNA-seq, but with a polyA-restricted library. If the 5’ usage bias is still apparent it suggests that pre-mRNAs, already decorated with polyA tails, depend on NONO for proper splicing, supporting post-transcriptional splicing.

How the formation of NONO RNA processing condensates impacts transcription initiation and elongation condensates remains open to debate. One possibility is that NONO condensates help release paused PolII (Adelman & Lis, 2012, Core, Waterfall et al., 2008, Lee, Li et al., 2008), or prevent promoter-proximal premature transcriptional termination (Ehrensberger, Kelly et al., 2013, Kamieniarz-Gdula & Proudfoot, 2019). Indeed, NONO co-purifies with some subunits of the mediator complex (required for release of paused PolII) (Huang, Li et al., 2012). A signature of paused PolII is abundant 20-60nt nascent RNA at the TSS (Rasmussen & Lis, 1993, Rougvie & Lis, 1988). Whilst we do see some genes with increased 5’ usage after NONO KD exhibiting such 20-60nt fragments, we also observe more extensive 5’ transcript increases (over the first 13% of transcript length). Increased abundance of reads at the TSS may also be an increase in nascent transcription of NONO-dependent genes to compensate for lower target protein levels. Additionally, NONO condensates may enhance PolII elongation, as SE’s generally regulate transcriptional elongation (Henriques, Scruggs et al., 2018, Tang, Yang et al., 2020) and SFPQ influences elongation (Hosokawa et al., 2019, Iida et al., 2020, Stagsted et al., 2021, Takeuchi et al., 2018). However, NONO KD leads to a significant upregulation only over the first 13% of the transcript, rather than a gradual decrease of transcript abundance across the entire gene. Further, PolII-Ser2 ChIP at the GATA2 locus is not altered by NONO KD. Thus whilst NONO condensates may influence PolII release and elongation to some extent, these are not the only mechanisms at play here.

NONO disruption leads to decreased expression of HAND2 and GATA2, transcription factors that are amongst a small group that maintain noradrenergic cell fate and survival in neuroblastoma (Boeva et al., 2017, Chipumuro et al., 2014, Durbin et al., 2018, Gartlgruber et al., 2021, Oldridge, Wood et al., 2015, van Groningen et al., 2017). High expression levels of these and other transcription factors work cooperatively in a defined core regulatory circuit creating a feed-forward loop to mediate sympathetic neuron specification, proliferation and differentiation (Rohrer, 2011, Voth et al., 2009, Willett & Greene, 2011). Enhancing expression of such targets suggests NONO may be part of a master transcriptional and post-transcriptional nexus that concertedly assemble a high density of transcription factors and coactivators to drive robust expression in neuroblastoma, which is in agreement with cancer cells that acquire SEs to drive expression of prominent oncogenes (Chapuy, McKeown et al., 2013, Hnisz, Shrinivas et al., 2017, Lovén, Hoke et al., 2013a). Thus NONO joins the ranks of oncogenic neuroblastoma coactivators such as BRD4, which forms nuclear phase-separated condensates at sites of SE-driven transcription (Sabari et al., 2018). Interestingly, BRD4/bromodomain inhibitors impair growth and induce apoptosis in neuroblastoma, hinting that potentially targeting NONO activity may also have therapeutic potential in this context (Puissant, Frumm et al., 2013). Furthermore, small molecules that interfere with RNA binding activity of NONO may serve as novel inhibitors of NONO biological functions and impede tumour cell growth. More broadly, this study may provide a new avenue for developing pharmaceutical drugs to manipulate RNA binding capacity of key regulators in combating cancer and other diseases.

## Materials and methods

### Cell culture and transfections

Two neuroblastoma cell lines including KELLY and BE(2)-C as well as HeLa cell line were used in this study. KELLY cells were grown in Gibco RPMI 1640 (ThermoFisher, 11835055) with 10% Serana foetal bovine serum (FBS) (Fisher Biotec, FBS-AU-015) and 100U/mL penicillin-streptomycin (PenStrep) (ThermoFisher, 15140122), whereas BE(2)-C and HeLa cell lines were grown in Gibco DMEM medium (ThermoFisher, 11995073) with 10% FBS and 100U/mL PenStrep. All cells were cultured in a 37°C incubator supplied with 5% CO_2_. These adherent cells were trypsinised using Gibco Tryple Express (ThermoFisher, 12604021) for routine passaging.

All transfections were performed in a forward transfection manner, with KELLY cells grown in RPMI (4% FBS) and HeLa cells grown in DMEM (10% FBS) during transfection. Lipofectamine 3000 (ThermoFisher, L3000015) and Lipofectamine RNAiMAX (ThermoFisher, 13778150) were the transfection reagents used in this study. All transfection mixtures were made up in serum-reduced Gibco Optimem (ThermoFisher, 11058021), as per the manufacturer’s instructions. The cells were collected 2 days after the final transfection unless otherwise stated. The siRNAs used in the study were silencer select negative control No. 1 siRNA (Thermofisher 4390844), Silencer Select siRNA NONO s9612 (Thermofisher 4392422) and Silencer Select siRNA NONO s9613 (Thermofisher, 4392420). Plasmid pEYFP-NONO containing siRNA resistance site corresponding to siRNA NONO s9612 was made by QuikChange site-directed mutagenesis kit (Stratagene, 200518) with base-pair substitutions and the template plasmid pEYFP-C1-NONO (human). Using siRNA resistant pEYFP-NONO plasmid as a template, plasmids pEYFP-NONO_ΔRRM1 (67-141 deletion), R256I and F257I were made by Q5 site-directed mutagenesis kit (New England BioLabs, E0554S) with base-pair substitutions according to the manufacturer’s instructions.

For siRNA-only transfections, 25 nM of siRNAs and Lipofectamine RNAiMAX were used. For pEYFP-control and pEYFP-NONO plasmids-only transfections, Lipofectamine 3000 was used, with 1.25 ug transfected into 6-wells for Western blots and 200 ng (HeLa cells) or 400 ng (KELLY cells) transfected into 12-wells for microscopic analysis. Sequential transfections for NONO RIP experiments were performed in 6-well plates using 10 nM of siRNAs and Lipofectamine RNAiMAX to reduce the endogenous NONO followed by 1.25 ug of plasmid DNA and Lipofectamine 3000 on the next day. Co-transfections of siRNAs and plasmids for BrdU assays were conducted in 12-well plates using Lipofectamine RNAiMAX with 25 nM of siRNAs and 1 ug of plasmid DNAs.

### RNA extraction and real time RT-qPCR

Cells were lysed with Nucleozol (Macherey-Nagel, 740404) and RNA extractions were conducted according to the manufacturer’s instructions. Lysed samples were heated at 55°C and vortexed at 1,000 rpm for 10 min. GlycoBlue coprecipitant (ThermoFisher, AM9516) was added prior to isopropanol precipitation to aid pellet visualisation.

RNA samples were reverse transcribed to cDNA using the QuantiTect reverse transcription kit (Qiagen, 205314). Real time qPCRs were performed in the Rotor-Gene Q real-time PCR cycler (Qiagen). A PCR reaction consisted of SensiMix SYBR No-ROX (Bioline, QT650-20), 250nM of forward and reverse primers (Integrated DNA Technologies, Table S1), molecular-grade H_2_O, and cDNA. Ribosomal protein P0 (RPLP0) and U6 spliceosomal RNA (U6) were used as reference genes and mRNA expression level relative to U6 was presented using 2-ΔΔCt method. To assess differential processing at the 5’ end of the transcripts, primer pairs over adjacent exon-intron junction (HAND2_2 and GATA2_2) were used to measure levels of pre-mRNAs, and primers pairs across exon-exon junction (HAND2_3) or located only in the exons (GATA2_3) were used to analyse the mature spliced transcripts. The level of pre-mRNA processing was then normalized over the expression of each mature spliced transcript. In ChIP-qPCR experiments, primer pairs of HAND2 and GATA2_4 were used to analyse the enrichment of chromatin fragments against RNA polymerase II phosphorated at Serine 2 with RPLP2 and CEP55 as positive and negative controls respectively. Primer pairs of GATA2_S, GATA2_M and GATA2_L were used to analyse the enrichment of chromatin fragments against NONO with GATA2_intron (equivalent to pre_GATA2) as a negative control.

### Protein extraction and Western blot

Cells were lysed with RIPA lysis buffer (150 mM NaCl, 25 mM Tris pH 7.5, 1% sodium deoxycholate, 0.1% SDS, 1% IGEPAL CA-630). Protein samples were mixed with SDS gel-loading buffer and heated to 95°C for 10 min. Samples and Precision Plus Protein™ All Blue Prestained Protein Standards (Bio-Rad, 1610373) were loaded onto Mini-PROTEAN TGX Pre-Cast Gels (Bio-Rad, 4561086). Gels were run in Tris/Glycine Buffer (Bio-Rad, 1610771) at 200V. Trans-Blot Turbo Mini Nitrocellulose Transfer packs (Bio-Rad, 1704158) were used for membrane transfer. Primary antibodies used were NONO (mouse monoclonal, in-house made), GATA2 (CG2-96) (mouse monoclonal, Santa Cruz, sc-267), HAND2 (A-12) (mouse monoclonal, Santa Cruz, sc-398167) and SFPQ (mouse monoclonal, Merck, P2860). Primary antibodies of NONO and SFPQ were diluted 1:1,000 in 5% milk PBST, while GATA2 and HAND2 were used at 1:500 dilution in PBST. The secondary (horseradish-peroxidase conjugated) antibodies including goat anti-mouse IgG H&L HRP (Abcam, ab97023) and goat anti-rabbit IgG H&L HRP (Abcam, ab97051) were diluted 1:10,000 in 5% milk PBST for NONO and SFPQ and 1:5,000 for GATA2 and HAND2. Luminata Crescedo Western HRP substrate (Merck, WBLUR0100) was added and blot images were acquired by Bio-Rad Chemidoc. Bio-Rad Imagelab Version 5.2 was used to quantify total protein levels and the intensity of the protein chemiluminescent bands. The relative intensity of chemiluminescent bands were normalized to the amount of total protein in each sample lane, and the sizes of chemiluminescent bands determined in relation to the standards ladder bands.

### GFP-trap

Cells (2.5 × 10^6^) were rinsed twice with ice-cold PBS and lysed in 150 uL RIPA buffer (25 mM Tris pH 7.5, 150 mM NaCl, 1% IGEPAL CA-630, 1% Sodium deoxycholate, 0.1% SDS) with 1 × protease inhibitor cocktail (Merck, 04693132001). The cell lysates were centrifuged at 20,000 g for 15 min at 4°C and resuspended in TN buffer (25 mM Tris pH 7.5, 150 mM NaCl, 0.5% IGEPAL CA-630). After taking pre-GFP-trap aliquots, the lysates were incubated with 10 μL GFP-Trap Magnetic Agarose beads (ChromoTek, gtma-20) for 3 h at 4°C, followed by washing three times with TN buffer. The samples were mixed with SDS buffer for Western blot.

### Immunofluorescence and RNA fluorescence in situ hybridization

Cells were grown on and fixed onto coverslips (Schott, G405-15) using 4% paraformaldehyde in PBS. Immunofluorescence started with permeabilisation in freshly made 1% Triton X-100 diluted in PBS for 10 min. Epitope detection was conducted with a primary monoclonal mouse antibody against NONO at 1:500 dilution in PBST and a polyclonal rabbit antibody against RNA Pol II phosphorated at Serine 2 (Abcam, ab5095) at 1:500 dilution. After three times of PBST washes, coverslips were incubated with secondary FITC anti-mouse antibody for NONO (Jackson Laboratories, 115-095-072) and TRITC anti-rabbit antibody for Pol II pSer2 (Jackson Laboratories, 711-026-152) at 1:500 dilution in PBST. Coverslips were then counter stained with DAPI and images acquired. For dual immunofluorescence, the cells were incubated with primary antibodies raised in different species together.

For dual immunofluorescence and RNA FISH, after immunofluorescence with NONO, coverslips were then hybridised overnight with FISH probes at 37°C according to the manufacturer’s instructions. The RNA FISH probes used in this study included human NEAT1 middle segment with Quasar R 570 Dye (Stellaris, SMF-2037-1) and custom-made human GATA2 and HAND2 segments with Quasar R 570 Dye (Stellaris).

For the treatment with 1,6-hexanediol, 1,6-hexanediol (Merck, 240117-50G) dissolved in the culture medium was added to cells at room temperature for 5 min. Then, the cells were fixed with 4% paraformaldehyde in PBS, followed by immunofluorescence procedure. The cells with low levels of overexpression were imaged.

Coverslips with cells grown on were rinsed in PBS and reaction buffer (20mM Tris pH 7.5, 5mM MgCl_2_, 0.5mM EGTA, 1 × protease inhibitor cocktail). Cells were permeabilised with 0.1% Triton X-100 diluted in reaction buffer for 5 min at room temperature, followed by rinsing subsequently in reaction buffer and PBS. Coverslips were then incubated with RNase A (Merck, R4642) or DNase I (Worthington, LS006333) to achieve a final concentration of 100 μg/mL in nuclease buffer (5 mM MgCl_2_ in PBS) for 20 min at room temperature. After nuclease digestion, cells were fixed with 4% paraformaldehyde in PBS and continued with immunofluorescence procedure.

### BrdU cell replication assay

KELLY cells were plated onto coverslips in 12-well plates and transfected with plasmids for 4 days. 2 hours prior to fixation, medium was replaced with the culture medium containing 10 nM 5-Bromo-2′-deoxyuridine (BrdU) (Merck, B5002-100MG). Coverslips were then fixed, and cells were permeabilised in 1% Triton X-100 diluted in PBS for 10 min at room temperature. After washing in PBS, coverslips were incubated in 1 M HCl for 30 min, rinsed twice in PBS and then incubated in 0.1 M sodium borate for 30 min, followed by blocking in 5% goat serum in PBST. Immunofluorescent staining was carried out using Anti-BrdU antibody (Abcam, ab8955) at 1:500 dilution in PBST and then anti-mouse Cy5-conjugated secondary antibody (Jackson Laboratories, 115-175-072) at 1:300 dilution in PBST.

### Expression and purification of recombinant proteins *in vitro*

The pEGFP-NONO_WT or pEGFP-NONO_ΔRRM1 plasmid was transformed into competent Rosetta 2(DE3)pLysS Escherichia coli cells (Novagen, 71400) and plated on LB agar plates with selection for kanamycin and chloramphenicol. When the optical density of the culture reaches 0.6 - 0.8, the flasks were cooled on ice for 15 minutes before expression was induced with 0.5 mM isopropyl β-D-1-thiogalactopyranoside. After incubation for 16 hours, the cells were harvested by centrifugation at 4000 g. A single pellet from 500 mL of expression culture was gently resuspended on ice in 50 mL binding buffer (50 mM Tris, pH 7.5, 1 M KCl, 300 mM L-arginine, 25 mM imidazole, 5% glycerol) supplemented with 5 μL Benzonase nuclease (Sigma, E1014-25KU), 1 mM phenylmethanesulfonyl fluoride, 1X cOmplete™ protease inhibitor cocktail (Roche, 11873580001) and 1 mM MgCl_2_. The cells were lysed with an Emulsiflex C5 high-pressure homogeniser (Avestin). The lysate was clarified by centrifugation at 24,000 g and then filtered through a 0.22 μM syringe filter. The supernatant was applied to a 5-ml His-Trap HF column charged with NiCl_2_ (GE Healthcare, 17-5248-02). GFP-NONO_WT or GFP-NONO_ΔRRM1 was eluted at room temperature with an imidazole gradient (25-500 mM) using the binding buffer and elution buffer (50 mM Tris, pH 7.5, 1 M KCl, 300 mM L-arginine, 500 mM imidazole, 5% glycerol) over ten column volumes. The peak fractions were pooled and loaded onto a Superdex 200 HiLoad 16/60 preparative-grade column (GE Healthcare, 17-1069-01) in 5 mL injections and developed at room temperature with the size exclusion buffer (20 mM Tris, pH 7.5, 1 M KCl, 50 mM L-Proline, 300 mM L-arginine, 0.5 mM EDTA, 5% glycerol). Relevant peak fractions were pooled and concentrated with Amicon Ultra Centrifugal Filter Units 30 kDa MWCO (Merck Millipore, UFC903024) to the required concentration.

### *In vitro* phase separation assay

Exogenously expressed NONO_WT and NONO_ΔRRM1, both tagged at the N-terminus with monomeric eGFP, were concentrated to 250 µM in storage buffer (20 mM Tris, pH 7.5, 500 mM KCl, 50 mM L-proline, 300 mM L-arginine, 0.5 mM EDTA, 5% glycerol). Four two-fold serial dilutions of the proteins were made in storage buffer to obtain proteins with concentration, 125, 62.5, 31.3 and 15.6 µM. Proteins at these five concentrations were diluted 1 in 10 in appropriate dilution buffers to 25, 12.5, 6.25, 3.13 and 1.56 µM. The composition of the dilution buffer was 20 mM Tris, pH 7.5, 50 mM L-proline, 0.5 mM EDTA, 5% glycerol and with the KCl concentration adjusted so that the final KCl concentrations after dilution were 50, 100, 150, 300 and 500 mM as required. Twenty µL of each of the diluted proteins were pipetted into wells of a 384-well flat-bottomed non-protein binding microplate (Grenier, #781906). The wells were sealed with clear film and incubated at room temperature for about 30 min. After incubation, DIC and FITC images of the wells were acquired with the Nikon Eclipse Ti2-E inverted microscope. To observe the effect of 2’-O-methyl phosphorothioate antisense oligonucleotides (PS-ASO) against NEAT1 on the phase separation of GFP-NONO_WT and GFP-NONO_ΔRRM1, the phase separation assays were repeated by diluting 1 in 10 the protein at 250 µM in appropriate dilution buffers to reach a final concentration of 100 mM KCL, 30 mM L-Arg and 0, 2, 4, 6, 8 and 10 µM PS-ASO after dilution, keeping other buffer component concentrations the same as the storage buffer.

### Microscopy and image analysis

All images were acquired on a Deltavision Elite microscope (GE) using a 60X for BrdU assays or 100X objectives for others. For subsequent counting and quantitative analysis of fluorescent intensities, the Nikon NiS Elements software (Version 4.3, Nikon, Tokyo, Japan) was used. Acquisition parameters were kept consistent and intensity thresholds were set the same for samples within each experiment. Cells that had incorporated BrdU into their DNA during replication (labelled by Cy5) were measured and expressed relative to the total number of nuclei, as measured by DAPI staining. MeanGreen of NONO within each RNA FISH foci or each immunofluorescent foci of other proteins was calculated as a ratio relative to nuclear MeanGreen of NONO, and then averaged for each cell. The ratio close to 1 indicates that NONO MeanGreen within specific foci is similar to nuclear NONO MeanGreen, implying no NONO enrichment and thus no co-localization between NONO signal and other molecules. Sphericity analysis was performed in ImageJ/Fiji with additional plugins (Haase, Jain et al., 2020, Haase, Royer et al., 2020, Legland, Arganda-Carreras et al., 2016, Ollion, Cochennec et al., 2013). Briefly, images were processed with CLIJ2 and the CLIJx-Assistant, NONO puncta were segmented using MorpholibJ’s Marker-controlled Watershed, and sphericity was calculated from the resulting objects in the 3D ImageJ Suite. All image processing and segmentation parameters were standardised between experimental groups.

### RNA immunoprecipitation (RIP)

KELLY cells (2.5 × 10^6^) were rinsed twice with ice-cold PBS, UV crosslinked at 400 mJ/cm^2^ on ice for 10 min in PBS, and then resuspended in 100 uL RIPA buffer (25 mM Tris pH 7.5, 150 mM NaCl, 1% IGEPAL CA-630, 1% Sodium deoxycholate, 0.1% SDS, 1 × protease inhibitor cocktail (Merck, 04693132001) and 2 U SUPERase-In RNase inhibitor (Thermofisher, AM2696)). Cells were sonicated using S220 Focused-ultrasonicator (Covaris, MA, USA) and cell lysates were centrifuged at 14,000 rpm for 15 min at 4 °C. The supernatants were diluted in 300 uL TN buffer with RNase inhibitor (25 mM Tris pH 7.5, 150 mM NaCl, 0.5% IGEPAL CA-630) and pre-cleared with 15 μL Dynabeads Protein G (Thermofisher, 10004D) for 3 h at 4°C. After taking pre-RIP aliquots, the pre-cleared supernatants were divided into two parts equally and incubated with 15 μL Dynabeads pre-bound with antibodies for 1.5 μg NONO or normal mouse IgG (Santa Cruz, sc-2025) overnight at 4 °C. The bead complexes were washed twice with RIPA buffer and twice with TN buffer. Then, the samples were incubated with TN buffer with 0.5% SDS and proteinase K (ThermoFisher, EO0491) at 55°C for 30 min, followed by RNA extraction. After reverse transcription, qPCR was used to amplify immunoprecipitated transcripts and data were presented as fold change relative to the input.

### Chromatin immunoprecipitation

KELLY cells (2 × 10^7^) were fixed in 10 mL PBS with 1% formaldehyde for 20 min at room temperature. Crosslinking was quenched by the addition of glycine to a final concentration of 0.125 M for 5 min. Cells were centrifuged at 3,000 rpm for 5 min and resuspended in lysis buffer (50 mM Hepes pH 7.5, 140 mM NaCl, 1 mM EDTA pH 8.0, 10% glycerol, 0.5% IGEPAL CA-630, 0.25% Triton X-100) with 1 × protease inhibitor cocktail, followed by rinsing twice in wash buffer (10 mM Tris pH 8.0, 200 mM NaCl, 1 mM EDTA pH 8.0, 0.5 mM EGTA). The cell pellets were resuspended in nuclei lysis buffer (50 mM Tris pH 8.0, 10 mM EDTA pH 8.0, 1% SDS) with 1 × protease inhibitor cocktail and sonicated with S220 Focused-ultrasonicator to achieve 200-500 bp DNA fragments. After centrifuging at 10,000 g for 10 min at 4 °C, the supernatants were mixed with ChIP dilution buffer (50 mM Tris pH 8.0, 167 mM NaCl, 1.1% Triton X-100, 0.11% sodium deoxycholate) and RIPA-150 buffer (50 mM Tris pH 8.0, 150 mM NaCl, 1 mM EDTA pH 8.0, 0.1% SDS, 1% Triton X-100, 0.1% sodium deoxycholate). The samples were pre-cleared with 20 μL Dynabeads and pre-ChIP aliquots taken. The pre-cleared supernatants were divided into two parts and incubated with 40 μL Dynabeads pre-bound with antibodies for 3.5 μg NONO or RNA polymerase II phosphorated at Serine 2 (Abcam, ab5095) and normal mouse IgG overnight at 4 °C. The bead complexes were sequentially washed twice each with RIPA-150 buffer, RIPA-500 buffer (50 mM Tris pH 8.0, 500 mM NaCl, 1 mM EDTA pH 8.0, 0.1% SDS, 1% Triton X-100, 0.1% sodium deoxycholate), RIPA-LiCl buffer (50 mM Tris pH 8.0, 1 mM EDTA pH 8.0, 1% IGEPAL CA-630, 0.1% sodium deoxycholate, 500 mM LiCl) and TE/10 buffer (10 mM Tris pH 8.0, 0.1 mM EDTA pH 8.0). The complexes were eluted by proteinase K in proteinase K digestion buffer (20 mM Hepes pH 7.5, 1 mM EDTA pH 8.0, 0.5% SDS) for 15 min at 50°C, followed by adding 3 μL 5 M NaCl and 1 μL 30 mg/mL RNase A for reverse crosslinking overnight at 65°C. The samples were further digested by proteinase K for 1 h at 50°C. The DNAs were then purified by SparQ Pure Mag beads (Quantabio, 95196-005) according to the manufacturer’s instructions and diluted in TE/10 buffer. qPCR was used to amplify immunoprecipitated chromatin fragments and data were presented as the percent input.

KELLY cells were treated with 100 μM 5,6-Dichlorobenzimidazole 1-β-D-ribofuranoside (DRB, Merck, D1916-10MG) for 3 h at 37°C. Cells in washout group were treated with DRB for 3 h at 37°C, followed by replacing with the normal culture medium for 2 h at 37°C. Cells from control, DRB and washout groups were harvested for ChIP-qPCR.

### RNA sequencing

KELLY cells were plated in 6-well plates and transfected with control siRNA and NONO KD siRNAs at a final concentration of 10 nM using Lipofectamine RNAiMAX for 72 hours. Cells (2.5 × 10^6^) were extracted for RNA samples as outlined above. RNA samples were sent to the Australian Genome Research Facility (AGRF) for sequencing. RNA quality was confirmed using a Bioanalyser (Perkin Elmar, MA, USA) prior to RNA-seq. Whole transcriptome libraries were prepared with the TruSeq stranded total RNA library prep kit (Illumina, CA, USA) and ribosomal RNA depleted with the Ribo-Zero-Gold kit (Illumina, CA, USA). Sequencing was performed using a HiSeq 2000 (Illumina, CA, USA) to generate 50 bp single end reads, resulting in an average 17-19 million reads per lane per sample. Reads from two lanes were pooled for each sample to generate 34-38 million reads for each sample.

### Phosphoactivatable ribonucleoside-enhanced crosslinking and immunoprecipitation

KELLY cells were grown on 20 x 14 cm dishes and treated for 14-16 hours with 100 µM of 4-Thiouridine (Merck, T4509). Cells (1.6 × 10^8^) were rinsed with ice-cold PBS, irradiated at 0.15 J/cm^2^ with 365nm UV light, scraped and resuspended in NP40 lysis buffer (50mM Hepes pH7.5, 1.5M KCl, 2mM EDTA, 0.5% NP-40, 0.5mM DTT, 1 × protease inhibitor cocktail). Then, cells were lysed and treated with 1 U/μL RNase T1 (Thermo Fisher Scientific, EN0541) for 5 min at room temperature to ensure that only RNA molecules bound by proteins were left. The lysates were pre-cleared with 20 μL Dynabeads for 30min at 4°C and then incubated with 100 μL Dynabeads conjugated with 50 μg NONO antibody for 2 h at 4°C. Samples were rinsed three times with NP40 lysis buffer and treated with 0.5 U/μL calf intestinal alkaline phosphatase (New England BioLabs, M0290) in dephosphorylation buffer (50 mM Tris pH 7.5, 100 mM NaCl, 10mM MgCl_2_, 1mM DTT) for 10 min at 37°C. The bead complexes were rinsed twice in phosphatase wash buffer (50 mM Tris pH 7.5, 20 mM EGTA, 0.5% NP40) and twice in polynucleotide kinase buffer (PNK, 50 mM Tris pH 7.5, 50 mM NaCl, 10 mM MgCl_2_). The samples were end-labelled with radioactive γ-^32^P-ATP (Perkin Elmar, NEG002250UC) to a final concentration of 0.5 μCi/μL and 0.8 U/μL T4 polynucleotide kinase (New England BioLabs, M0201S) for 30 min at 37°C. Then, the bead complexes were rinsed five times in PNK buffer and resuspended in 2 × SDS buffer. After SDS-PAGE electrophoresis, gels were visualized on films and gel bands containing the target crosslinked protein-RNA complexes cut out. Gel bands were treated to electro-elution and the eluted complexes in SDS buffer were further incubated with 1% SDS and proteinase K for 30 min at 55°C. RNA extraction was then carried out using the miRNeasy Micro Kit (Qiagen, 217084) and RNA samples were sequenced.

### Bioinformatics

Raw sequencing files were quality checked using FastQC (version 0.11.5) with all files passing. All subsequent analysis was performed using the gencodev37lift37 transcript model.

#### PAR-CLIP

PARpipe (https://github.com/ohlerlab/PARpipe) was used with the default parameters to process PAR-CLIP datasets. In brief, after pre-processing, the pipeline uses PARalyzer (version 1.5) to identify reads containing T-to-C transitions (which are indicative of successful RNA-protein crosslinking), and aggregates these reads into clusters to define RNA-protein binding sites. Alignment files from PARpipe were analysed using wavClusteR (version 2.24.0) to determine and annotate binding sites. TPM was calculated as described (Wagner et al., 2012) and used to define NONO-bound genes as those in the top 20^th^ percentile of genes featuring reads with T-to-C transitions. Gene ontology analysis was performed for NONO bound genes using Metascape (http://metascape.org, (Tripathi, Pohl et al., 2015)), with expressed genes as the background. PAR-CLIP coverage plots were generated using metagene2 (version 1.6.1).

#### Gene-level RNA-seq

Transcript quantification was performed using salmon (version 1.4.0), summarised to gene-level counts and imported into R using tximport (1.18.0). Differential expression analysis was performed using DESeq2 (1.30.1) and the default parameters (alpha = 0.1). In the NONO KD siRNA experiment, four control samples were tested against eight NONO KD samples. Samples from the two NONO KD siRNAs were grouped in an attempt to correct for off-target effects of the individual siRNAs. Gene ontology analysis was performed for differentially expressed genes using Metascape (http://metascape.org, (Tripathi et al., 2015)), separating genes by positive or negative log fold change (LFC). Gene Set Enrichment Analysis (GSEA, version 3.0) was also performed using the count data generated above and tested against the Biological Process gene sets maintained at MSigDB (https://software.broadinstitute.org/gsea/msigdb/).

#### Splicing analysis

IsoformSwitchAnalyzeR (version 1.12.0) was run using output from salmon to look for overall splicing changes. To extend this, DEXSeq (version 1.36.0) was used to extract all exons from the gencodev37lift37 gtf, and custom scripts were used to extract all introns. DESeq2 and DEXSeq were then run to determine differential expression and differential usage respectively, at the level of exons and introns (separately). Then, for each gene, the maximum length transcript was divided into 100 bins. The position of each significant differentially expressed and differentially used exon and intron was then overlapped with these 100 bins. The number and direction of change of significant events within each bin was summed. Bins for which the proportion of positive fold change events was 2 standard deviations greater than the median proportion were deemed as significant. Heatmaps were created for all genes which showed significant events within the first 13 bins (as defined by significant difference in the proportion of positive and negative events). Heatmaps show the log2 fold change of significant events as a function of their relative position within the gene.

#### Super enhancer detection from ChIP-seq

Publicly available KELLY (Chipumuro et al., 2014) cell line ChIP-seq data for the H3K27ac histone mark (GSM1532401) and associated input (GSM1532403) were downloaded from GEO. ChIP-seq reads were aligned using Bowtie 2 (Langmead & Salzberg, 2012) against the hg19 (GRCh37) reference genome with the parameters -k 2 -q. Aligned reads were filtered, sorted and indexed using SAMtools (Li, Handsaker et al., 2009) using a minimum mapping quality score of 30. ChIP-seq peak calling was performed using MACS2 (Zhang, Liu et al., 2008) with the parameters: --keep-dup auto –p 1e-9 -B. Super enhancer calling was performed using ROSE (Lovén, Hoke et al., 2013b, Whyte, Orlando et al., 2013) with the parameters: -s 12500 -t 1000. Promoter associated enhancer exclusion in ROSE was changed to a 1000bp window (+/-500bp) to allow the detection of MYCN associated super enhancer regions. Annotation of associated gene TSS for super enhancer regions was done using the org.Hs.eg.db and TxDb.Hsapiens.UCSC.hg19.knownGene packages in R. All genes whose TSS fell within 600kbp flanking of detected super enhancer start and end sites were included as associated genes.

#### Survival Analysis

The Kaplain-Meier tool from R2 genomics (https://hgserver1.amc.nl/cgi-bin/r2/main.cgi?option=kaplan_main) was used to identify neuroblastoma datasets which showed MYCN as having a prognostic value for overall survival using the ‘scan’ method and default settings. Using these four datasets (Cangelossi, Maris, SEQC and Versteeg) the tool was then used to identify all genes which were of prognostic value, using the ‘median’ method and default settings. The median method here was used as a more conservative means of identifying prognostic genes of otherwise unproven clinical relevance. Any gene whose expression was significantly associated with prognosis in at least 3 of the 4 datasets was then taken as showing prognostic value in neuroblastoma.

## Acknowledgements

We thank other members of the Fox lab for helpful discussions about the manuscript. This work was supported by a research grant APP1147496 from the National Health and Medical Research Council of Australia to AF and CB and FT180100204 from the Australian Research Council to AF.

## Author contributions

SZ performed experiments and analysed data. JC analysed RNA sequencing data. YSC, AN, TL, SA, SK, OM and YSC performed experiments. SZ, JC and AF wrote the manuscript and prepared the figures. All authors read and approved the manuscript.

## Tables and their legends

**Table S1.**
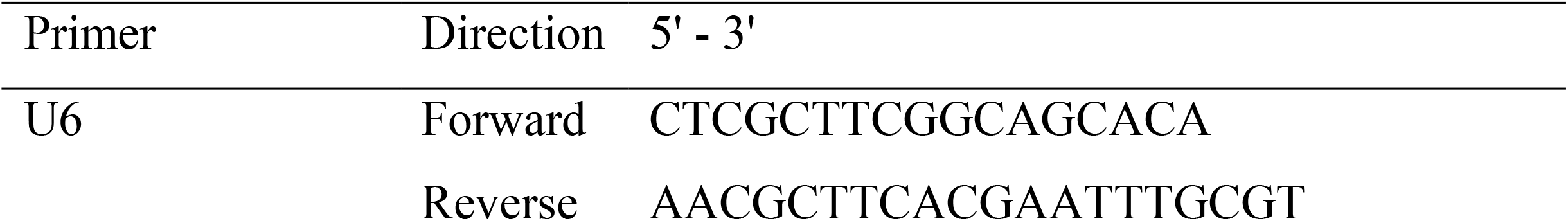
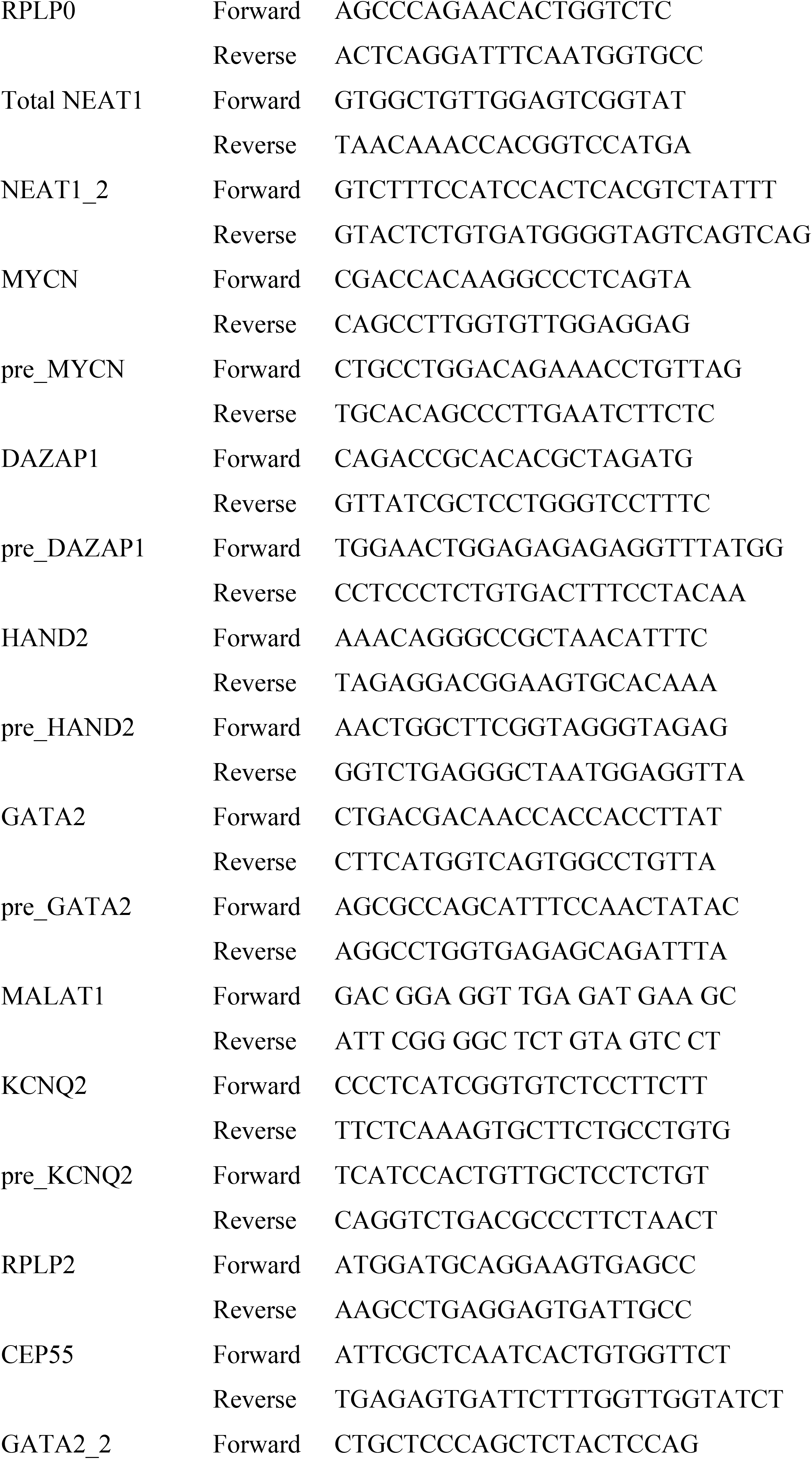
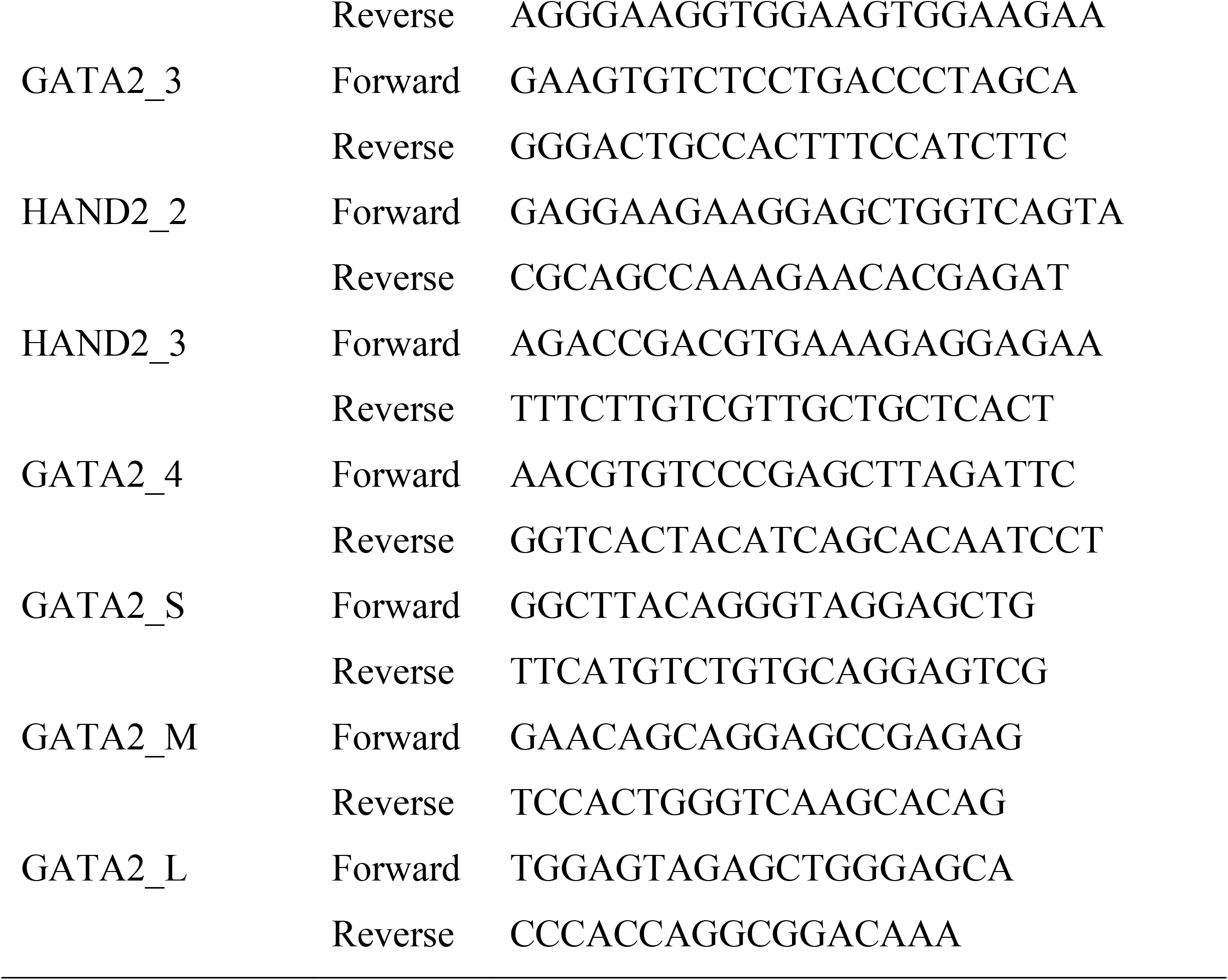
A list of primers.

## Expanded View Figure legends

**Supplementary figure 1:**
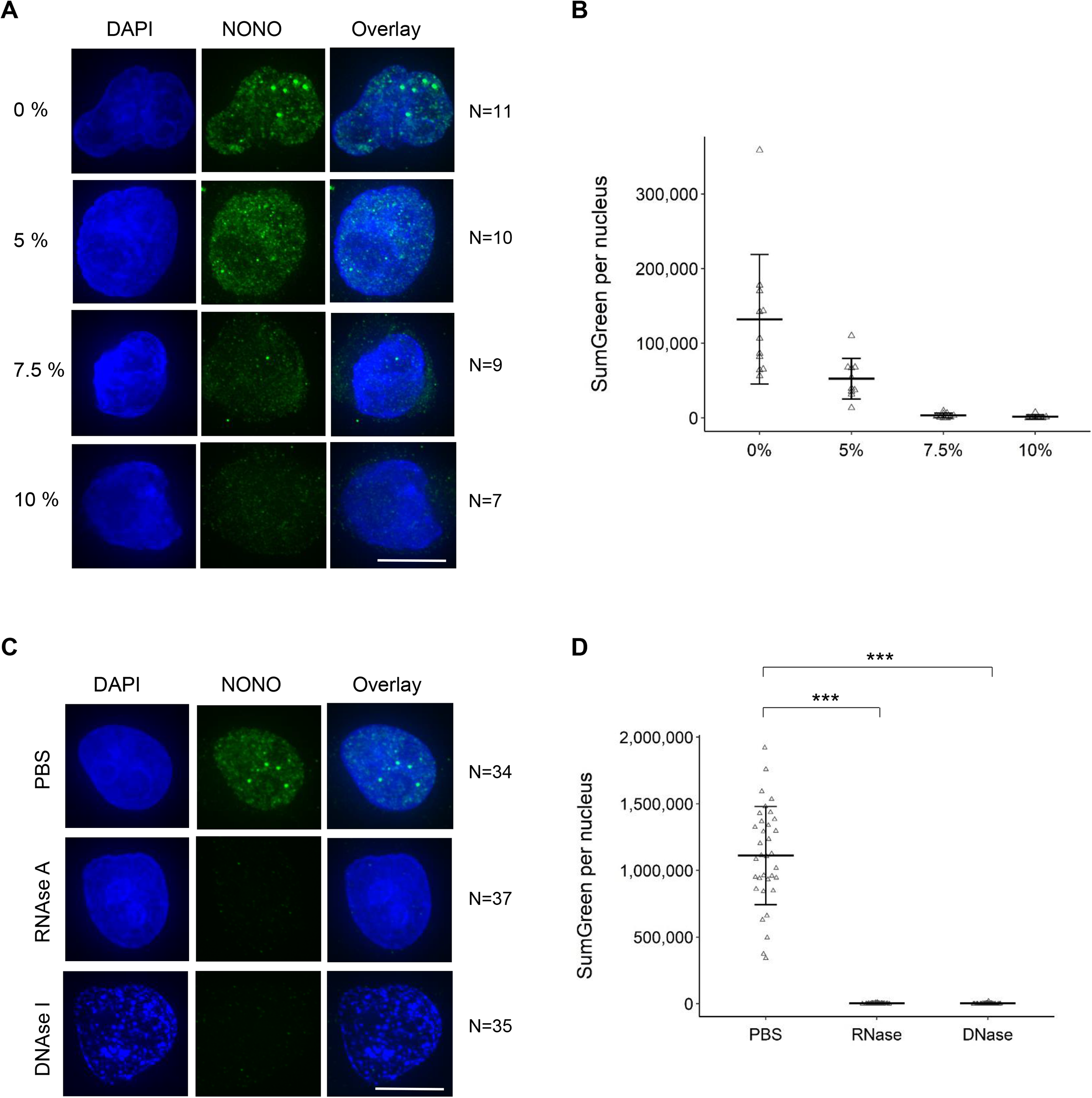
Both RNA and DNA are required for NONO distribution in Hela cells. (A) Fluorescence micrograph images of representative cells treated with 0, 5, 7.5 or 10% 1,6 hexanediol. DAPI (blue) stain indicates cell nuclei and NONO immunofluorescence (green). (B) Dot plot of SumGreen per nucleus at different concentrations of 1,6 hexanediol in (A). Bars are SD. (C) Fluorescence micrograph images of representative cells treated with PBS, RNase A or DNase I and stained for NONO. (D) Dot plot of SumGreen per nucleus in (C). Bars are SD. ***P<0.001.

**Supplementary figure 2:**
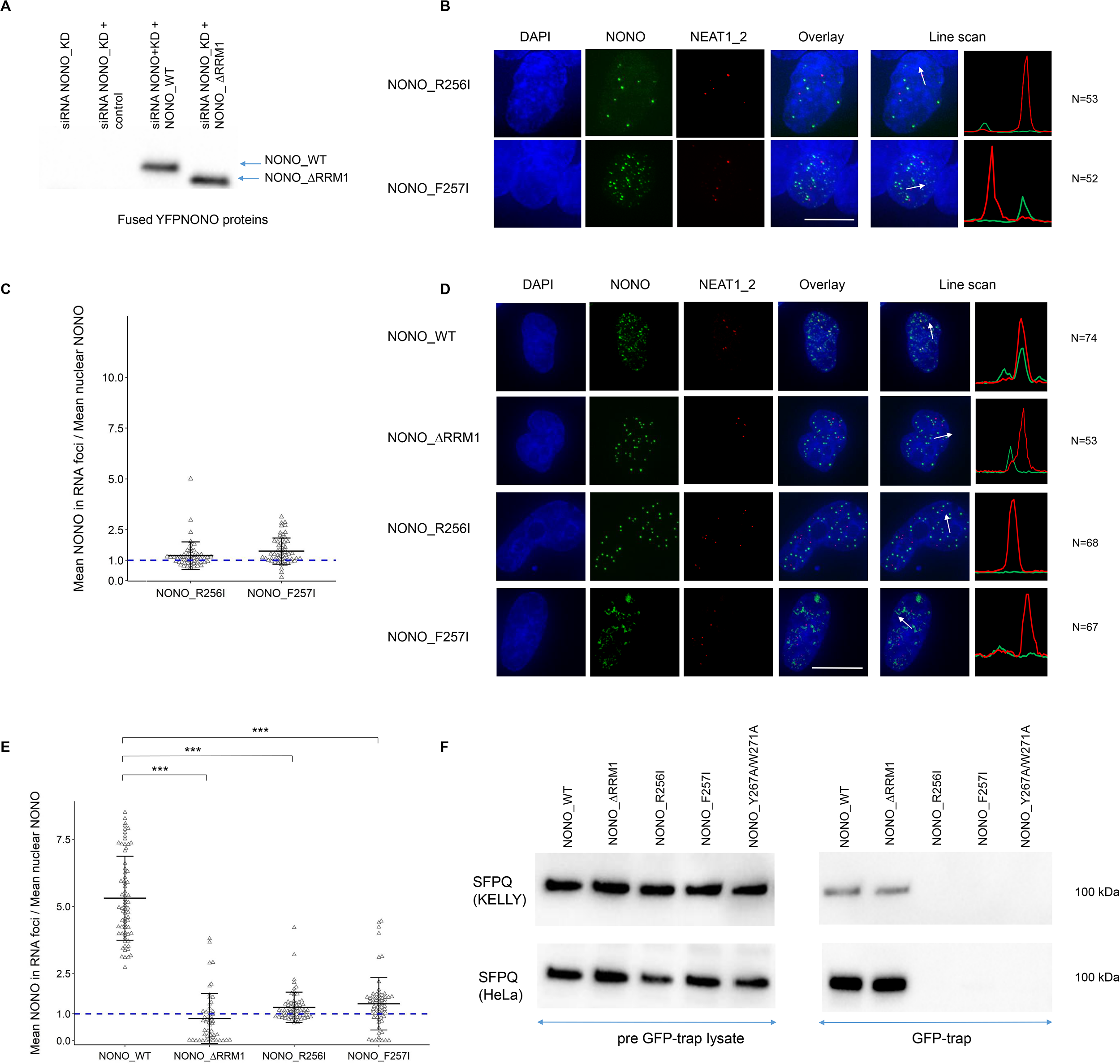
Two single mutants of NONO play roles in dimerization. (A) Representative Western blot images of YFP fused NONO_WT and NONO_ΔRRM1 proteins in KELLY cells. (B) Fluorescence micrograph images of representative KELLY cells stained for NONO and NEAT1_2 after transfection with YFP fused NONO_R256I and NONO_F257I exogenous protein plasmids. DAPI (blue) stain indicates cell nuclei, YFP fused NONO (green) and NEAT1_2 RNA FISH (red). Scale bar: 5μm. (C) The enrichment of mean NONO fluorescence detected within RNA FISH foci is quantitatively determined as a ratio relative to mean nuclear NONO fluorescence in (B). Bars are SD. (D) Fluorescence micrograph images of representative HeLa cells stained for NONO and NEAT1_2 after transfection with YFP fused NONO_WT, NONO_ΔRRM1, NONO_R256I and NONO_F257I exogenous protein plasmids. DAPI (blue) stain indicates cell nuclei, YFP fused NONO (green) and NEAT1_2 RNA FISH (red). Scale bar: 5μm. (E) The enrichment of mean NONO fluorescence detected within RNA FISH foci is quantitatively determined as a ratio relative to mean nuclear NONO fluorescence in (D). Bars are SD. ***p<0.001. (F) Representative Western blot images of SFPQ protein in pre-GFP-trap lysate samples and GFP-trapped samples transfected with YFP fused NONO_WT, NONO_ΔRRM1, NONO_R256I, NONO_F257I and NONO_Y267A/W271A exogenous protein plasmids in KELLY and HeLa cells.

**Supplementary figure 3:**
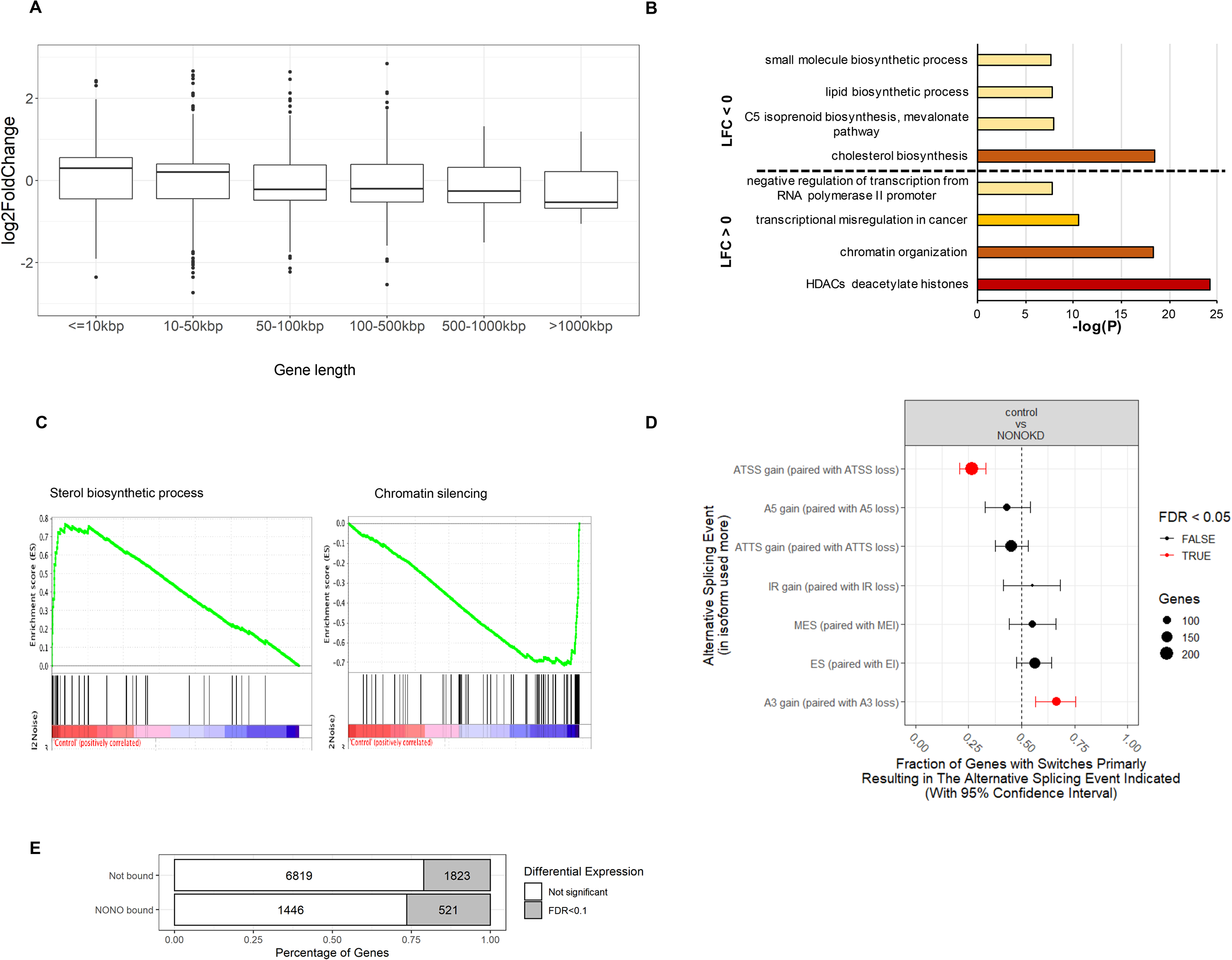
NONO KD induces significant changes in ATSS and A3 in KELLY cells. (A) Box plots showing log fold change for genes which were significantly differentially expressed (padj < 0.1), separated by gene length. (B) Summary of most highly enriched gene ontology categories produced by ‘Metascape’ analysis of significantly upregulated (LFC > 0) and downregulated (LFC<0) genes following NONO KD. (C) Enrichment score profile plots produced by ‘GSEA’ for the ‘sterol biosynthetic process’ and ‘chromatin silencing’ gene sets respectively. (D) NONO KD results in significant splicing events including ATSS and A3. (E) Proportions of differentially expressed genes between NONO bound and non-bound target genes.

**Supplementary figure 4:**
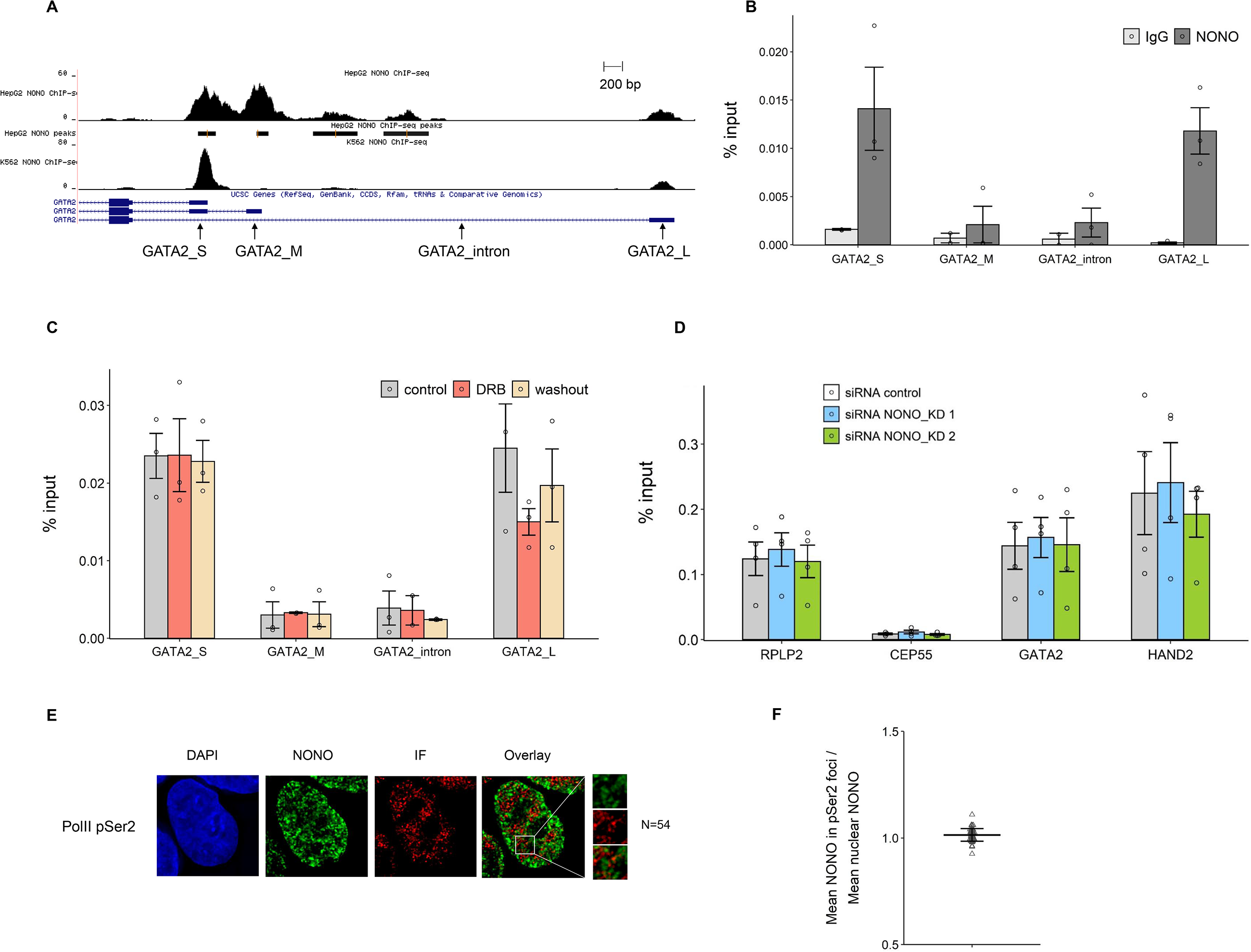
NONO function is dependent on DNA binding, which in turn is not sensitive to transcription levels. (A) NONO ChIP-seq peaks for GATA2 in HepG2 and K562 cell lines shown as a Genome Browser image (Xiao et al., 2019). The primer pairs for GATA2 used in our ChIP-qPCR are indicated. (B) The enrichment of relative GATA2 chromatin fragments via ChIP-qPCR against NONO and normal mouse serum IgG in KELLY cells. Bars are SEM. n≥3. (C) NONO ChIP-qPCR indicates the enrichment of relative GATA2 chromatin fragments in KELLY cells treated with control, DRB and DRB followed by a washout period. Bars are SEM. n≥3. (D) The enrichment of relative GATA2 and HAND2 chromatin fragments after NONO KD via ChIP-qPCR against RNA PolII phosphorated at Serine 2 in KELLY cells. Bars are SEM. n≥3. (E) Fluorescence micrograph images of representative cells stained for NONO and RNA PolII phosphorated at Serine 2. DAPI (blue) stain indicates cell nuclei, NONO immunofluorescence (green) and immunofluorescence (red) for Pol II pSer2. Scale bar: 5μm. (F) In micrograph image quantitation analysis, the enrichment of mean NONO fluorescence detected within immunofluorescence foci of Pol II pSer2 is determined as a ratio relative to mean nuclear NONO fluorescence in (E). Bars are SD.

